# Sequencing RNA from old, dried specimens reveals past viromes and properties of long-surviving RNA

**DOI:** 10.1101/2024.10.03.616531

**Authors:** Alexandra H. Keene, Mark D. Stenglein

**Affiliations:** Center for Vector-Borne and Infectious Diseases, Department of Microbiology, Immunology, and Pathology, College of Veterinary Medicine and Biomedical Sciences, Colorado State University, Fort Collins, CO, USA; Quantitative Cell and Molecular Biology Graduate Program

## Abstract

Recovery of virus sequences from old samples provides an opportunity to study virus evolution and reconstruct historic virus-host interactions. Studies of old virus sequences have mainly relied on DNA or on RNA from fixed or frozen samples. The millions of specimens in natural history museums represent a potential treasure trove of old virus sequences, but it is not clear how well RNA survives in old samples. We experimentally assessed the stability of RNA in insects stored dry at room temperature over 72 weeks. Although RNA molecules grew fragmented, RNA yields remained surprisingly constant. RT-qPCR of host and virus RNA showed minimal differences between dried and frozen specimens. To assess RNA survival in much older samples we acquired *Drosophila* specimens from North American entomological collections. We recovered sequences from known and novel viruses including several coding complete virus genomes from a fly collected in 1908. We found that the virome of *D. melanogaster* has changed little over the past century. Galbut virus, the most prevalent virus infection in contemporary *D. melanogaster*, was also the most common in historic samples. Finally, we investigated the genomic and physical features of surviving RNA. RNA that survived was fragmented, chemically damaged, and preferentially double stranded or contained in ribonucleoprotein complexes. This showed that RNA - especially certain types of RNA – can survive in biological specimens over extended periods in the absence of fixation or freezing and confirms the utility of dried specimens to provide a clearer understanding of virus evolution.

## IMPORTANCE

RNA from old specimens has been instrumental for understanding the origin and evolution of RNA viruses. Most studies have relied on relatively rare fixed or frozen samples, likely because researchers assume that RNA doesn’t survive well. Using experimentally dried insects and dried specimens from museums, we show that RNA can in fact persist for decades or longer without freezing or fixation. We found that the virome of the model fruit fly *Drosophila melanogaster* has remained largely stable over the last century. But we also discovered completely new viruses from past infections. Double-stranded RNA preferentially survived, but double strandedness did not account for all surviving RNA, and RNA incorporated into proteinaceous complexes also survived better. This work confirms the value of the millions of dried specimens stored in natural history and archaeological collections for understanding the evolutionary history of RNA viruses.

## INTRODUCTION

The study of ancient DNA has been a useful way to better understand the evolutionary history of cellular organisms and their pathogens^1–4^. RNA viruses cause many important diseases, but the study of old RNA has mainly relied on relatively rare fixed or frozen specimens^5,6^. Nevertheless, such specimens have been useful for reconstructing the early history of pandemic pathogens^7,8^. Other studies have taken advantage of endogenized RNA virus sequences, which capture the sequences of viruses that existed long ago ^9–11^. The understanding of virus evolution would benefit from the recovery of additional old RNA virus sequences. But how well RNA survives in the absence of freezing or fixation is unclear.

There are reasons to believe that RNA may not survive well over extended periods. For one thing, RNA is short-lived during normal cellular function. The half-life of messenger and ribosomal RNAs are measured in minutes or hours in a living cell^12^. RNA molecules are shorter than genomic DNA and more prone to hydrolysis at physiological conditions^13^. Marketing of RNA-stabilizing reagents further reinforces the idea that RNA is unstable. Studies of historic RNA may therefore be uncommon simply because researchers assume that RNA does not survive well in old specimens.

However, there are an increasing number of studies that indicate that RNA may in fact survive well over long periods^13^. Examples include recovery of a single stranded (ss) RNA virus genome from 750-year-old barley, sequencing of a double stranded (ds) RNA virus genome from 1000-year-old corn, and profiling of RNA from a 130-year-old extinct Tasmanian tiger^14–16^. These studies indicate that RNA can survive for long periods and hint at the potential broad utility of old specimens to study RNA viruses from hundreds or thousands of years ago.

There are nearly 300 million archived arthropod specimens in North American museums alone^17^. Dried beetles, butterflies, ants and fruit flies have yielded useful DNA but to our knowledge RNA has not been sampled from such specimens^18–24^. Amongst the diverse organisms contained in entomological collections are flies in the *Drosophilidae* family^24^*. Drosophila melanogaster* has been instrumental to our understanding of genetics, evolution, and disease, and the virome of contemporary *D. melanogaster* has been described in detail^25–29^.

In this study we used two approaches to investigate RNA survival. First, we experimentally dried and froze flies and mosquitoes and measured RNA yield, length, and detectability over 72 and 52 weeks. Next, we obtained *Drosophila* specimens from museum collections. Using next generation sequencing we characterized surviving viral and host RNA. We recovered viral genome sequences from over 100 years ago and identified properties of RNA associated with survival. We followed strict lab protocols to minimize contamination by contemporary nucleic acid and present many lines of evidence to support the idea that our conclusions are based on RNA from the actual samples. Our findings challenge the assumption that RNA does not survive well and confirm the usefulness of dried biological specimens as a source of RNA to study past viral infection.

## RESULTS

### Viral and host RNA persisted in dried biological specimens

Entomological collections contain millions of dried specimens, but it’s not clear to what extent RNA survives in dried insects, or more generally in dried biological specimens. To assess RNA survival in such samples, we recovered RNA from flies and mosquitoes that had been pinned and stored at room temperature or that had been frozen. We used adult *D. melanogaster* and *Aedes aegypti* from outbred colonies maintained in our insectary^30,31^. Both populations contained individuals persistently infected by one or more viruses. We sampled *D. melanogaster* over 72 weeks and *Ae. aegypti* over 52 weeks. We quantified the concentration and size distribution of recovered RNA and used RT-qPCR to quantify levels of specific host and viral RNAs.

RNA concentrations recovered from individual flies were variable but we were able to recover RNA from all dried samples (**Fig. 1A**). We used a multiple linear regression model (MLR) to assess the relationship between concentration, time and storage condition. There was a small but statistically significant decrease in RNA concentration over time (F = 4.9, p = 0.03) but surprisingly RNA yields from dried and frozen samples did not significantly differ (F = 0.84, p = 0.36). We assessed the size distribution of recovered RNA and found that RNA grew increasingly fragmented in both dried and frozen flies (**Fig. 1B, Supp.** Fig. 1). In a multiple linear regression model with the same predictors as above, there was a statistically significant decrease in RNA length over time (F = 19.8, p = 4.3x10^-5^) but no significant difference in length between dried and frozen samples (F = 0.4, p = 0.53).

**Figure 1:**
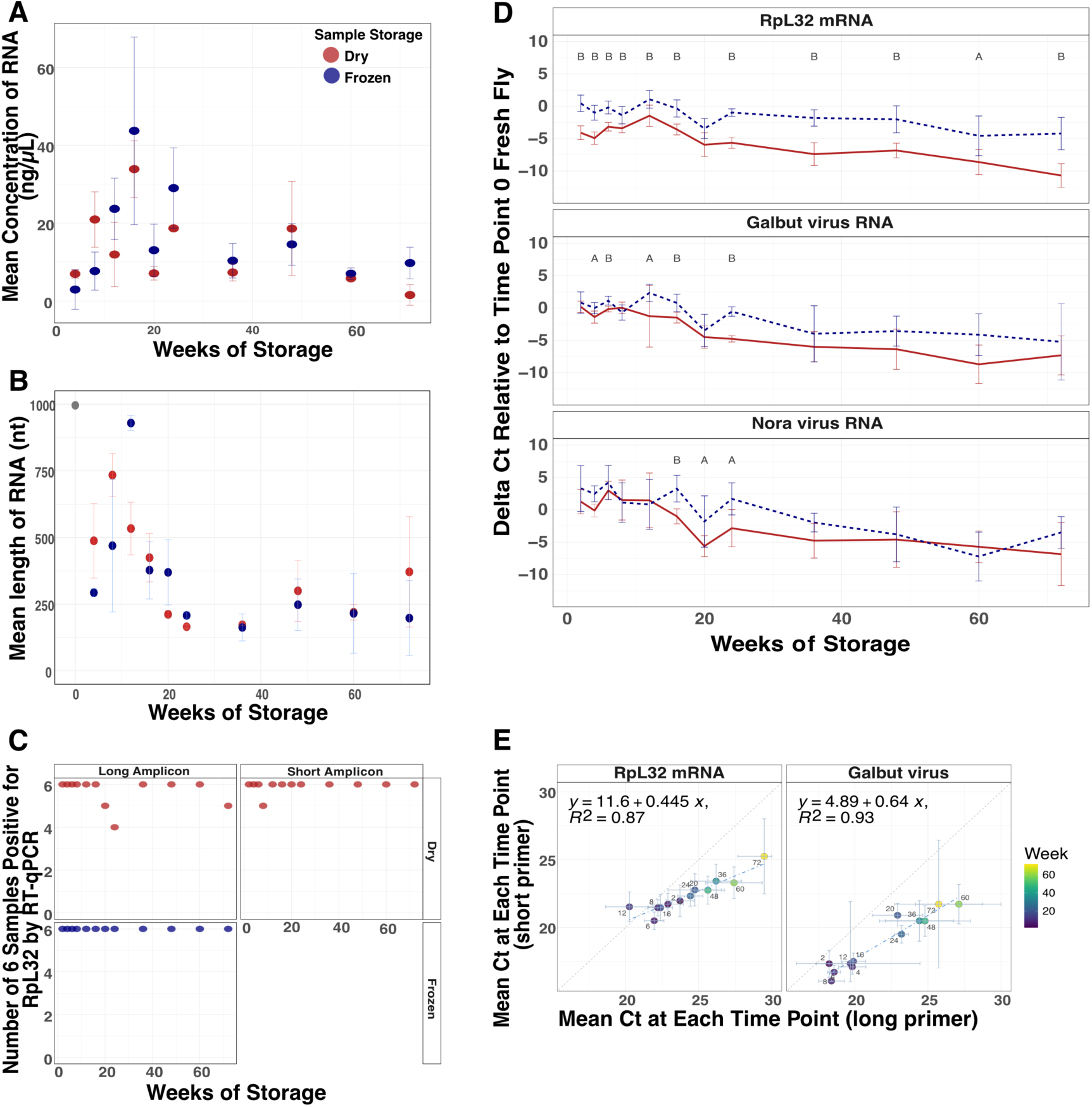
RNA survived in dried flies over 72 weeks. (A) Average RNA concentration and (B) length of RNA from dried and frozen flies over 72 weeks. N = 3 female *D. melanogaster* from each storage condition at each timepoint. (C) Number of samples positive of 3 male and 3 female flies at each timepoint for RpL32 mRNA via RT-qPCR using primers that target long (110 nt) and short (56 nt, dried only) regions of RpL32 mRNA. (D) Average difference of RpL32 mRNA, galbut virus and Nora virus RNA levels relative to fresh *D. melanogaster* collected at the initial timepoint. A two-tailed t-test was used to determine the statistical significance between dried and frozen means at every time point (A < 0.05, B < 0.001). (E) Comparison of mean Ct value at each time point of dried *D. melanogaster* using primers that target a long (x-axis) or short (y-axis) region of RpL32 mRNA and galbut virus. Error bars represent mean plus and minus the standard deviation. N = 3 male and 3 female at each time point.

Host messenger RNA (mRNA) remained reliably detectable by RT-qPCR in both dried and frozen samples. In *D. melanogaster*, we detected ribosomal protein L32 (RpL32) mRNA in 72 of 72 frozen flies and 68 of 72 dried flies (**Fig. 1C**). When we used shorter-range primers that amplified a 56 bp region of RpL32 instead of the 110 bp region amplified by our standard primers, 71 of 72 dried flies were positive (**Fig. 1C**).

Although there was no statistically significant difference in RNA concentration and length between dried and frozen flies, there was variability in detected levels of specific host and viral RNA targets (**Fig. 1D; Supp.** Fig. 2). Levels of host and viral RNA measured by RT-qPCR decreased as storage time increased. Levels of RpL32 mRNA were higher in frozen flies than they were in the dried flies (**Fig. 1D**).

Each qPCR run included one positive control: RNA from known infected flies, and three negative controls: an extraction blank (a no sample extraction), a no-sample RT control, and a no template qPCR control. The use of negative controls corresponding to each step of the extraction and detection process would have allowed us to determine when cross-contamination had been introduced if it had been. All controls behaved as expected, confirming the absence of cross-contamination during extraction and RT-qPCR (**Supp.** Fig. 3A).

The FoCo-17 *D. melanogaster* population we sampled contained individual flies variably infected by galbut virus, La Jolla virus, Nora virus, and Thika virus^32,33^. The prevalence of each virus varied in this outbred population as did viral RNA levels in infected individuals. As with RpL32 mRNA, average levels of detected viral RNAs decreased over time in both dried and frozen flies (**Fig. 1D; Supp.** Fig. 2A). Levels of galbut virus dropped by 69x in frozen samples and 181x in dried samples between 0 and 72 weeks. After 24 weeks, there was no significant difference between galbut virus RNA levels in dried and frozen flies. The overall pattern was similar for the three other viruses (**Fig. 1D; Supp Fig. 2**). Shorter range primers detected galbut virus RNA in fewer qPCR cycles, unsurprisingly confirming that short PCR products work better for fragmented RNA (**Fig. 1E**).

**Figure 2:**
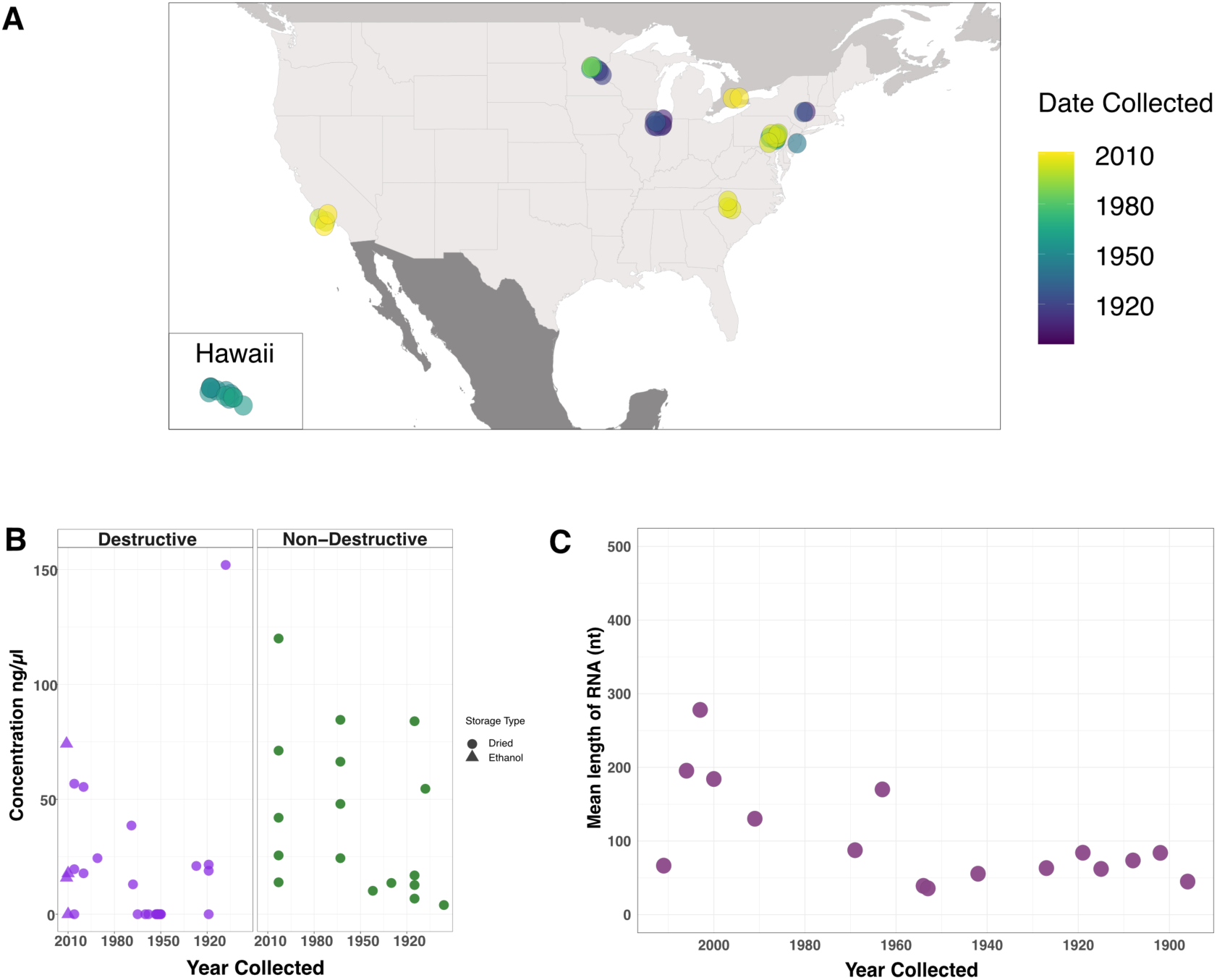
Museum sample collection locations and characteristics of extracted RNA. (A) Map of the United States and Canada indicating sample collection locations. (B) RNA concentration from individual museum flies using either the destructive (purple) or non-destructive (green) extraction method. (C) Mean length of RNA molecules from a subset of museum specimens encompassing the range of years specimens obtained for this study had been collected.

RNA also survived well in dried mosquitoes (**Supp.** Fig 4). An MLR was used to assess whether time and/or storage condition impacted mosquito RNA concentration. Over time (F = 2.09, p = 0.15) and between dried and frozen mosquitoes (F = 2.37, p = 0.13) there was no statistically significant difference in mosquito RNA concentrations (**Supp.** Fig. 4A). A caveat is that we neglected to save fresh mosquitoes at time 0 as we did with flies, so RNA levels were normalized to levels in the 4-week frozen samples, which had likely already experienced degradation. There was no statistically significant difference in RNA length over time after 4 weeks (F = 0.75, P = 0.39) but there was by storage condition (F = 79.3, p = 2.5x10^-13^; **Supp.** Fig. 4B).

We used RT-qPCR to assess levels and detectability of specific host and viral RNAs in stored mosquitoes. We detected actin mRNA in all 78 dried and all 78 frozen mosquitoes (**Supp.** Fig. 4C). Actin mRNA levels remained consistent over time with no difference between 4 weeks and 52 weeks in the frozen samples and a 6x decrease in the dried samples. This mosquito population harbored verdadero virus and Guadeloupe mosquito virus^30,34^. Viral RNAs decreased over time in the dried and frozen specimens. In the dried mosquitoes, virus levels dropped between 13x for verdadero virus and 18x for Guadeloupe mosquito virus between 4 and 52 weeks. In frozen mosquitoes there was no difference in verdadero virus levels between 4 weeks and 52 weeks but there was a 5x decrease for Guadeloupe mosquito virus (**Supp.** Fig. 4D-E).

The mosquito positive and negative controls behaved as expected except for the cDNA positive control sample at week 28, which was negative for Guadeloupe mosquito virus. This may reflect a true negative infection status, given that our cDNA positive control samples were individual mosquitoes from a variably infected outbred population confirmed to have a verdadero virus infection (**Supp.** Fig. 3B).

### *Drosophila* obtained for RNA metagenomic sequencing were 10-125 years old and from entomological collections across the United States and Canada

Having determined that RNA survives well in experimentally dried insects, we turned our attention to much older samples. We obtained 46 specimens from entomological collections across the United States and Canada (**Table 1**). Thirty-two had been assigned as *D. melanogaster* based on morphology, nine as *D. simulans*, and five as unknown drosophilids. Collection dates ranged from 1896 to 2011 (**Fig. 2A**; **Table 1; Supp. Table 1**). We extracted RNA from 29 specimens following a standard destructive protocol, which yielded RNA concentrations from 0 to 152 ng/µL (**Fig. 2B**; **Table 1; Supp. Table 1**). Some samples were loaned on the condition that they not be destroyed, so we followed a non-destructive RNA isolation protocol for the remaining samples^35^. This used an overnight incubation with proteinase to liberate internal contents of the specimen while leaving morphology intact. The non-destructive extractions yielded more RNA than the standard protocol (p = 0.03), with concentrations ranging from 4 to120 ng/µL (**Fig. 2B**; **Table 1; Supp. Table 1**). However, when samples from Hawai’i, which had unusually low yields, were removed there was no statistically significant difference between RNA concentration obtained from the two methods (p = 0.59; **Supp.** Fig. 5). Four samples that had been stored in 70% ethanol required rehydration prior to extraction (**Supp.** Fig 5).

**Table 1:**
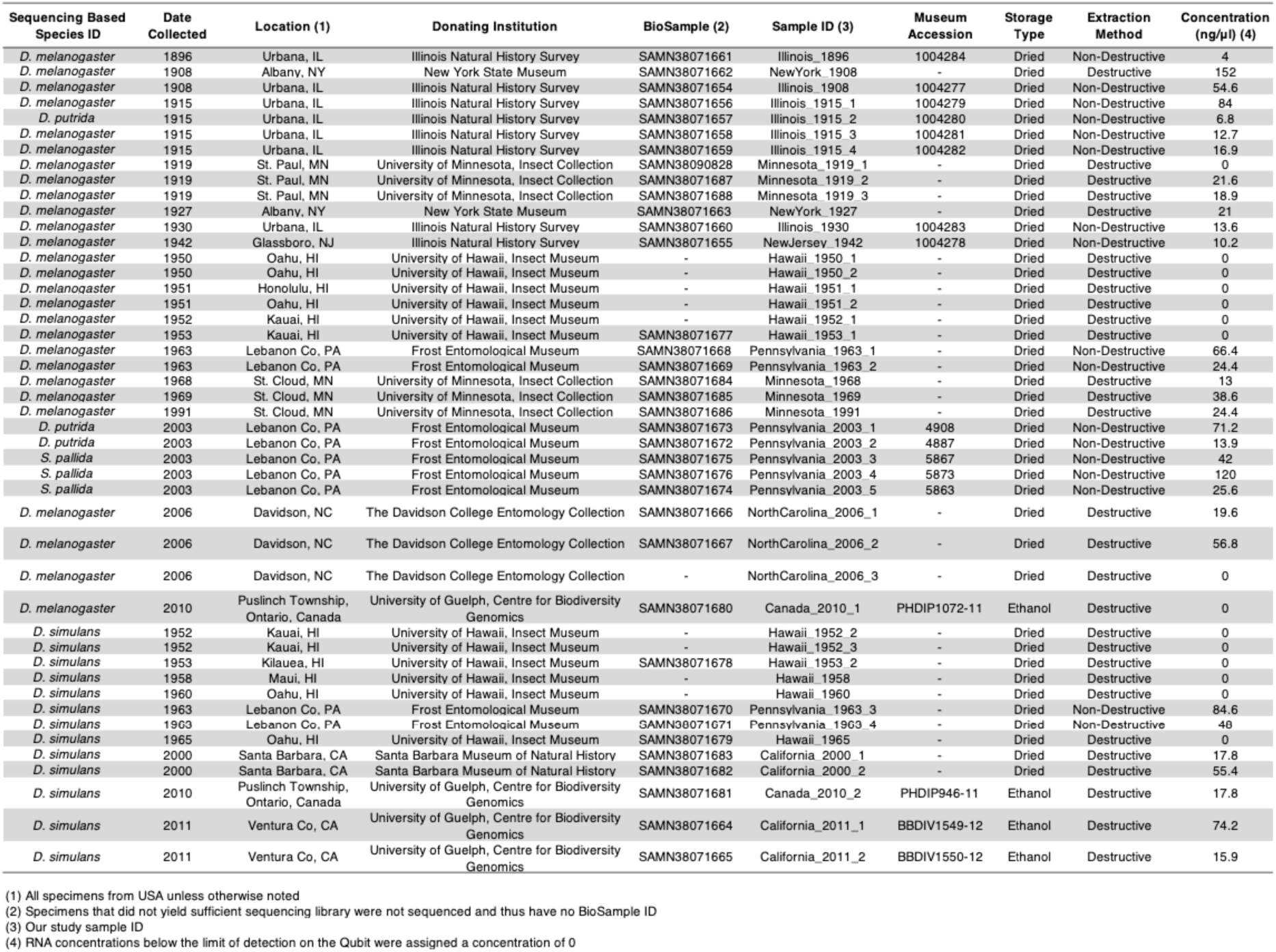
Museum specimen metadata.

RNA from samples collected after or before 1960 had average lengths of 159 and 60 nt, respectively (**Fig. 2C**). Although there was variability in RNA quantity and quality, the RNA obtained from these specimens was of suitable length and concentration for downstream analyses.

We screened museum samples for galbut virus RNA and RpL32 mRNA using our short-range primers. Sixteen of 46 samples (35%) were positive for galbut virus and 21 (46%) were positive for RpL32 mRNA (**Supp.** Fig. 6A**, Supp. Table 1**). Positive status was confirmed using melting temperature and agarose gel electrophoresis. The small size of PCR products made them difficult to distinguish from primer dimers on agarose gels, so it is possible that there were false negative calls. All negative control samples were negative as expected (**Supp.** Fig. 6B).

### Metagenomic sequencing of museum samples and molecular species assignment

We prepared shotgun sequencing libraries from museum sample RNA. Thirty-six samples yielded libraries suitable for sequencing. Sequencing yielded an average of 48 million reads per sample (range: 0.4-238 million). Several samples were sequenced multiple times to increase recovery of complete virus sequences.

The records for some museum samples did not include morphological taxonomic assignments below the family level, so we used competitive mapping to generate molecular taxonomic assignments. We mapped quality and adapter trimmed reads to a set of 545 representative cytochrome oxidase subunit-1 (CO1) sequences from across the Drosophilidae family to generate a molecular species identification for each sample. This molecular species assignment corroborated the museum morphology-based assignment for most samples (**Supp. Table 2)**. Four samples labeled as *D. melanogaster* in museum records were revealed to be *D. simulans* by CO1-mapping. *D. melanogaster* and *D. simulans* are common, sympatric, and difficult to distinguish morphologically, so such misassignment is expected^36^. Five samples did not have species-level taxonomic assignments from the museums. Of these, CO1-mapping indicated that 2 were *Drosophila putrida* and the remaining three were *Scaptomyza pallida*. One fly collected in Illinois in 1915 was also assigned as *D. putrida* (**Supp. Table 1-2**). These assignments are consistent with the known range of *D. putrida* and *S. pallida^37^*. All fresh frozen and experimental dried samples were assigned as *D. melanogaster* as expected and there were no CO1-mapping reads in the three water negative control datasets.

**Table 2:**
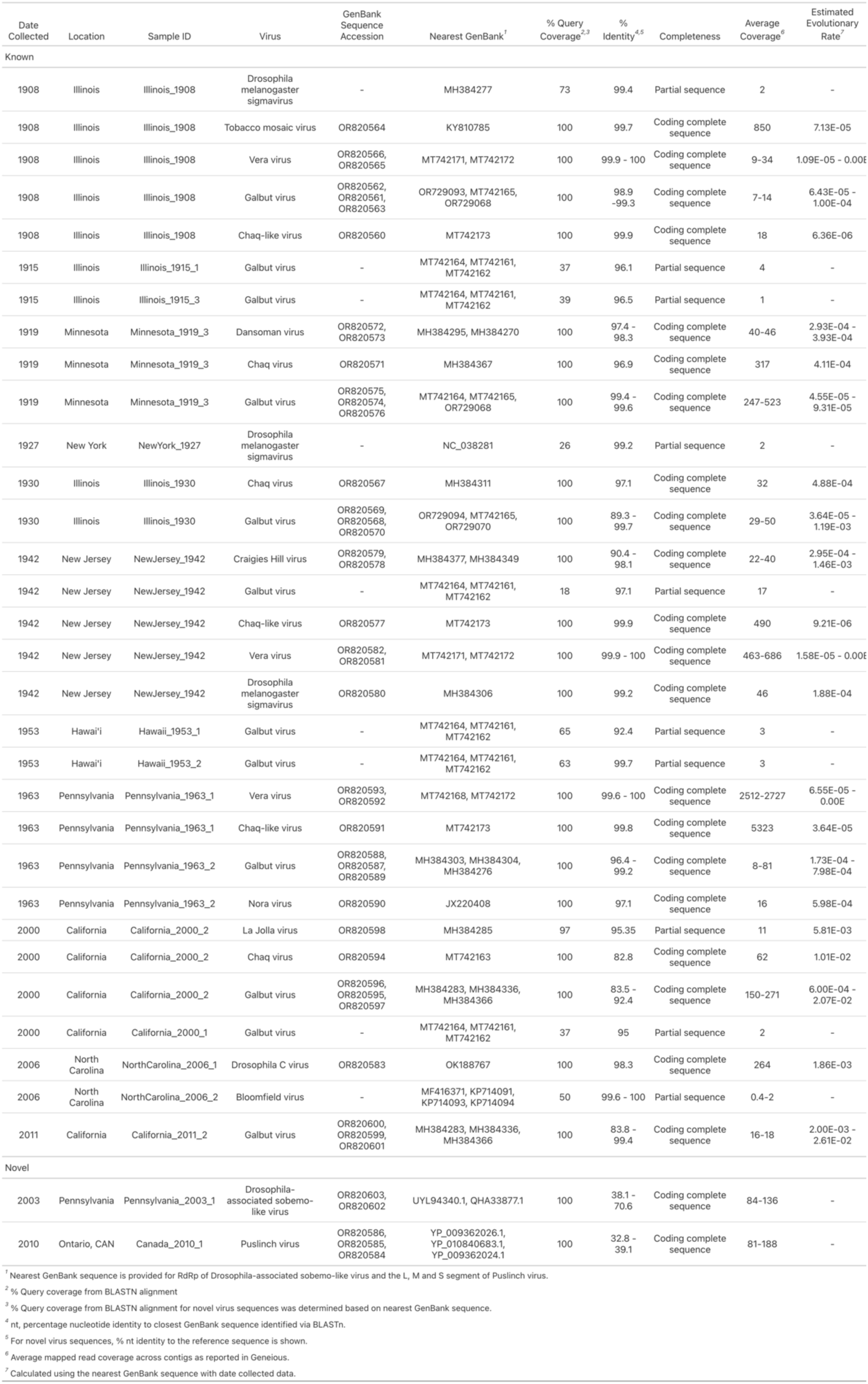
Virus sequences identified in *Drosophila* museum collection specimens.

| Date Collected | Location | Sample ID | Virus | GenBank Sequence Accession | Nearest GenBank<sup>1</sup> | % Query Coverage<sup>2,3</sup> | % Identity<sup>4,5</sup> | Completeness | Average Coverage<sup>6</sup> | Estimated Evolutionary Rate<sup>7</sup> |
| --- | --- | --- | --- | --- | --- | --- | --- | --- | --- | --- |
| Known |  |  |  |  |  |  |  |  |  |  |
| 1908 | Illinois | Illinois_1908 | Drosophila melanogaster sigmavirus | - | MH384277 | 73 | 99.4 | Partial sequence | 2 | - |
| 1908 | Illinois | Illinois_1908 | Tobacco mosaic virus | OR820564 | KY810785 | 100 | 99.7 | Coding complete sequence | 850 | 7.13E-05 |
| 1908 | Illinois | Illinois_1908 | Vera virus | OR820566, OR820565 | MT742171, MT742172 | 100 | 99.9 - 100 | Coding complete sequence | 9-34 | 1.09E-05 - 0.00E-05 |
| 1908 | Illinois | Illinois_1908 | Galbut virus | OR820562, OR820561, OR820563 | OR729093, MT742165, OR729068 | 100 | 98.9 - 99.3 | Coding complete sequence | 7-14 | 6.43E-05 - 1.00E-04 |
| 1908 | Illinois | Illinois_1908 | Chaq-like virus | OR820560 | MT742173 | 100 | 99.9 | Coding complete sequence | 18 | 6.36E-06 |
| 1915 | Illinois | Illinois_1915_1 | Galbut virus | - | MT742164, MT742161, MT742162 | 37 | 96.1 | Partial sequence | 4 | - |
| 1915 | Illinois | Illinois_1915_3 | Galbut virus | - | MT742164, MT742161, MT742162 | 39 | 96.5 | Partial sequence | 1 | - |
| 1919 | Minnesota | Minnesota_1919_3 | Dansoman virus | OR820572, OR820573 | MH384295, MH384270 | 100 | 97.4 - 98.3 | Coding complete sequence | 40-46 | 2.93E-04 - 3.93E-04 |
| 1919 | Minnesota | Minnesota_1919_3 | Chaq virus | OR820571 | MH384367 | 100 | 96.9 | Coding complete sequence | 317 | 4.11E-04 |
| 1919 | Minnesota | Minnesota_1919_3 | Galbut virus | OR820575, OR820574, OR820576 | MT742164, MT742165, OR729068 | 100 | 99.4 - 99.6 | Coding complete sequence | 247-523 | 4.55E-05 - 9.31E-05 |
| 1927 | New York | NewYork_1927 | Drosophila melanogaster sigmavirus | - | NC_038281 | 26 | 99.2 | Partial sequence | 2 | - |
| 1930 | Illinois | Illinois_1930 | Chaq virus | OR820567 | MH384311 | 100 | 97.1 | Coding complete sequence | 32 | 4.88E-04 |
| 1930 | Illinois | Illinois_1930 | Galbut virus | OR820569, OR820568, OR820570 | OR729094, MT742165, OR729070 | 100 | 89.3 - 99.7 | Coding complete sequence | 29-50 | 3.64E-05 - 1.19E-03 |
| 1942 | New Jersey | NewJersey_1942 | Craigies Hill virus | OR820579, OR820578 | MH384377, MH384349 | 100 | 90.4 - 98.1 | Coding complete sequence | 22-40 | 2.95E-04 - 1.46E-03 |
| 1942 | New Jersey | NewJersey_1942 | Galbut virus | - | MT742164, MT742161, MT742162 | 18 | 97.1 | Partial sequence | 17 | - |
| 1942 | New Jersey | NewJersey_1942 | Chaq-like virus | OR820577 | MT742173 | 100 | 99.9 | Coding complete sequence | 490 | 9.21E-06 |
| 1942 | New Jersey | NewJersey_1942 | Vera virus | OR820582, OR820581 | MT742171, MT742172 | 100 | 99.9 - 100 | Coding complete sequence | 463-686 | 1.58E-05 - 0.00E-05 |
| 1942 | New Jersey | NewJersey_1942 | Drosophila melanogaster sigmavirus | OR820580 | MH384306 | 100 | 99.2 | Coding complete sequence | 46 | 1.88E-04 |
| 1953 | Hawai'i | Hawaii_1953_1 | Galbut virus | - | MT742164, MT742161, MT742162 | 65 | 92.4 | Partial sequence | 3 | - |
| 1953 | Hawai'i | Hawaii_1953_2 | Galbut virus | - | MT742164, MT742161, MT742162 | 63 | 99.7 | Partial sequence | 3 | - |
| 1963 | Pennsylvania | Pennsylvania_1963_1 | Vera virus | OR820593, OR820592 | MT742168, MT742172 | 100 | 99.6 - 100 | Coding complete sequence | 2512-2727 | 6.55E-05 - 0.00E-05 |
| 1963 | Pennsylvania | Pennsylvania_1963_1 | Chaq-like virus | OR820591 | MT742173 | 100 | 99.8 | Coding complete sequence | 5323 | 3.64E-05 |
| 1963 | Pennsylvania | Pennsylvania_1963_2 | Galbut virus | OR820588, OR820587, OR820589 | MH384303, MH384304, MH384276 | 100 | 96.4 - 99.2 | Coding complete sequence | 8-81 | 1.73E-04 - 7.98E-04 |
| 1963 | Pennsylvania | Pennsylvania_1963_2 | Nora virus | OR820590 | JX220408 | 100 | 97.1 | Coding complete sequence | 16 | 5.98E-04 |
| 2000 | California | California_2000_2 | La Jolla virus | OR820598 | MH384285 | 97 | 95.35 | Partial sequence | 11 | 5.81E-03 |
| 2000 | California | California_2000_2 | Chaq virus | OR820594 | MT742163 | 100 | 82.8 | Coding complete sequence | 62 | 1.01E-02 |
| 2000 | California | California_2000_2 | Galbut virus | OR820596, OR820595, OR820597 | MH384283, MH384336, MH384366 | 100 | 83.5 - 92.4 | Coding complete sequence | 150-271 | 6.00E-04 - 2.07E-02 |
| 2000 | California | California_2000_1 | Galbut virus | - | MT742164, MT742161, MT742162 | 37 | 95 | Partial sequence | 2 | - |
| 2006 | North Carolina | NorthCarolina_2006_1 | Drosophila C virus | OR820583 | OK188767 | 100 | 98.3 | Coding complete sequence | 264 | 1.86E-03 |
| 2006 | North Carolina | NorthCarolina_2006_2 | Bloomfield virus | - | MF416371, KP714091, KP714093, KP714094 | 50 | 99.6 - 100 | Partial sequence | 0.4-2 | - |
| 2011 | California | California_2011_2 | Galbut virus | OR820600, OR820599, OR820601 | MH384283, MH384336, MH384366 | 100 | 83.8 - 99.4 | Coding complete sequence | 16-18 | 2.00E-03 - 2.61E-02 |
| Novel |  |  |  |  |  |  |  |  |  |  |
| 2003 | Pennsylvania | Pennsylvania_2003_1 | Drosophila-associated sobemo-like virus | OR820603, OR820602 | UYL94340.1, QHA33877.1 | 100 | 38.1 - 70.6 | Coding complete sequence | 84-136 | - |
| 2010 | Ontario, CAN | Canada_2010_1 | Puslinch virus | OR820586, OR820585, OR820584 | YP_009362026.1, YP_010840683.1, YP_009362024.1 | 100 | 32.8 - 39.1 | Coding complete sequence | 81-188 | - |

<sup>1</sup> Nearest GenBank sequence is provided for RdRp of Drosophila-associated sobemo-like virus and the L, M and S segment of Puslinch virus. <sup>2</sup> % Query coverage from BLASTN alignment <sup>3</sup> % Query coverage from BLASTN alignment for novel virus sequences was determined based on nearest GenBank sequence. <sup>4</sup> nt, percentage nucleotide identity to closest GenBank sequence identified via BLASTn. <sup>5</sup> For novel virus sequences, % nt identity to the reference sequence is shown. <sup>6</sup> Average mapped read coverage across contigs as reported in Geneious. <sup>7</sup> Calculated using the nearest GenBank sequence with date collected data.

### The overall taxonomic composition of new and old samples was similar

To broadly assess the source of sequences in our datasets, we taxonomically binned sequences using or lab’s metagenomic classification pipeline ^38^. This first identified host-derived reads by mapping to an index composed of the *D. melanogaster*, *D. putrida*, and *S. pallida* genomes^39,40^. The pipeline then assembled remaining non-host reads and taxonomically assigned the resulting contigs and unassembled reads using a BLASTN search of the NCBI nucleotide database. We tabulated the fraction of reads in each dataset that were assigned to various taxa or that remained unassigned (**Fig. 3**).

**Figure 3:**
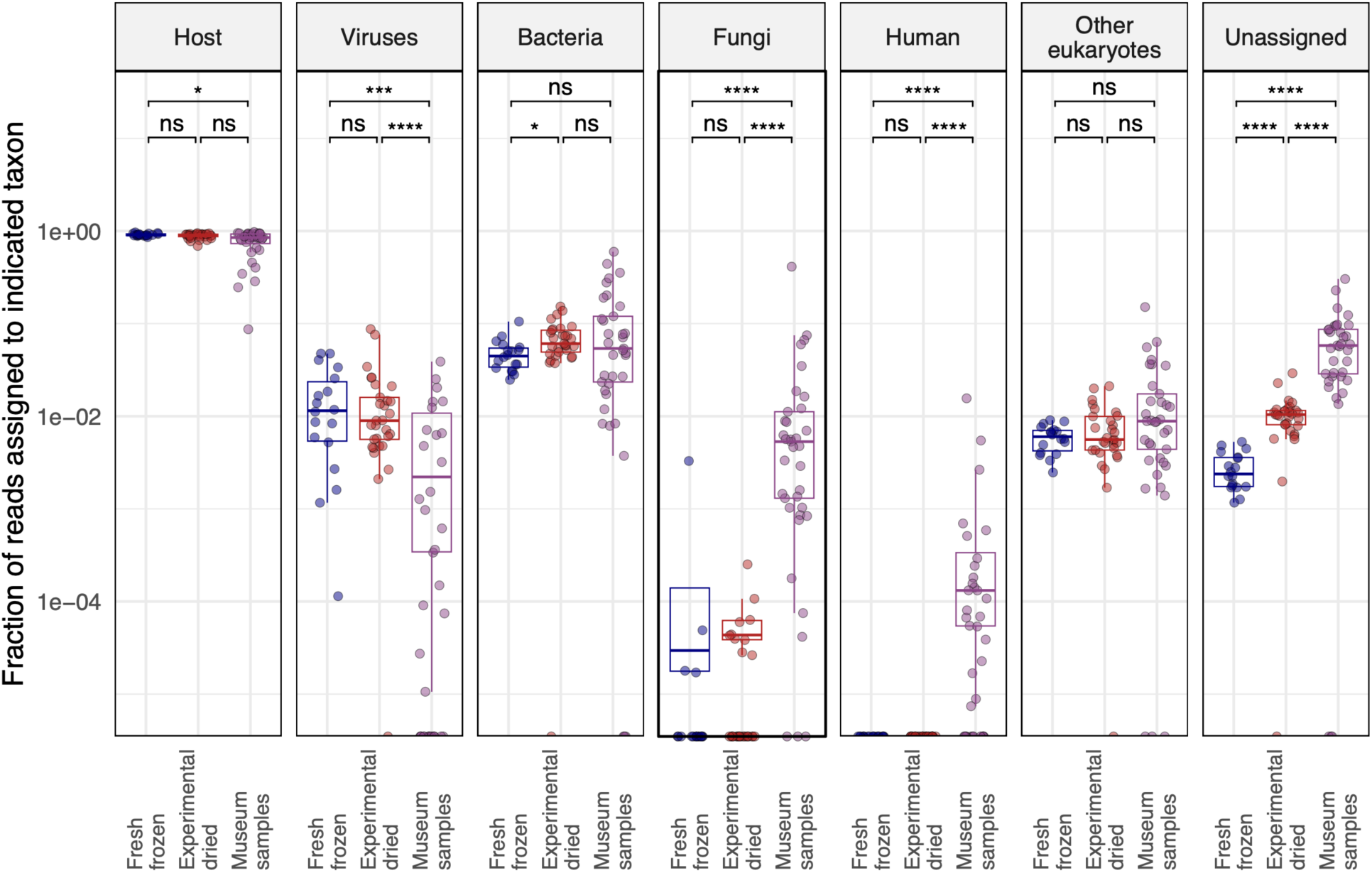
The overall taxonomic composition of old and new samples was similar. The fraction of reads assigned to the indicated taxa are plotted. Each point represents a dataset from an individual fly. Adjusted *p*-values significance levels from Wilcoxon test are indicated. Host-mapping reads were identified by mapping to combined *D. melanogaster*, *D. putrida*, and *S. pallida* genomes. Fractions of reads mapping to other taxa were determined by metagenomic classification of non-host-mapping reads. The “other eukaryotes” category accounts for all eukaryotic taxa apart from fungi and human. Unassigned read fractions account for reads that were not assigned by host-mapping or metagenomic classification.

Most reads mapped to the fly genomes in all datasets, although the median fraction of host-mapping reads fell from 90.5% in fresh samples to 85% in museum samples (**Fig. 3**). This decrease was accompanied by a corresponding increase in unassigned reads. Only 0.2% of reads were unassigned in fresh datasets, compared to 5.5% in museum datasets (p=2.3x10^-6^). This increase in unassignable reads is likely attributed to the fact that short reads from fragmented RNA contain less information for host mapping or taxonomic assignment. There were on average fewer virus-mapping reads in museum samples, which could reflect the fact that our FoCo-17 population was persistently infected by multiple viruses or could reflect decreased relative survival of viral RNA in old samples. There were increases in the proportion of fungi- and human-mapping reads in museum datasets. Fungal reads in old samples may have derived from saprophytic fungi. Human-mapping reads in museum datasets may have originated from handling of specimens by curators, accumulation of dust on samples, or an increased fraction of contamination-derived sequences in old datasets. The median fraction of human-mapping reads in old datasets was 5.4x10^-5^ and the maximum fraction was 1.6% of reads, in a dataset from Hawai’i. We concluded that some of the reads in our libraries, including human-assigned reads, may have derived from contamination. However, these accounted for a small fraction of datasets and the overall taxonomic composition of new and old datasets was similar (**Fig. 3**).

### Entomological specimens harbored diverse viral sequences including previously unknown viruses

We recovered at least one virus sequence from 21 of the 36 sequenced museum samples and two or more virus sequences from 5 samples (**Fig. 4**; **Table 2**). All but three of the virus sequences corresponded to known *Drosophila*-infecting viruses^25,26^. Galbut virus was the most common virus, with six samples producing coding-complete genomes and six samples producing partial galbut virus genomes (**Fig. 4**; **Table 2**). The next most common viruses were *D. melanogaster* sigma virus and vera virus, which were detected in three samples each (**Fig. 4**; **Table 2**). The sample with the most viral sequences was collected in 1908 in Illinois, USA: this fly yielded three coding-complete and one partial virus sequence.

**Figure 4.**
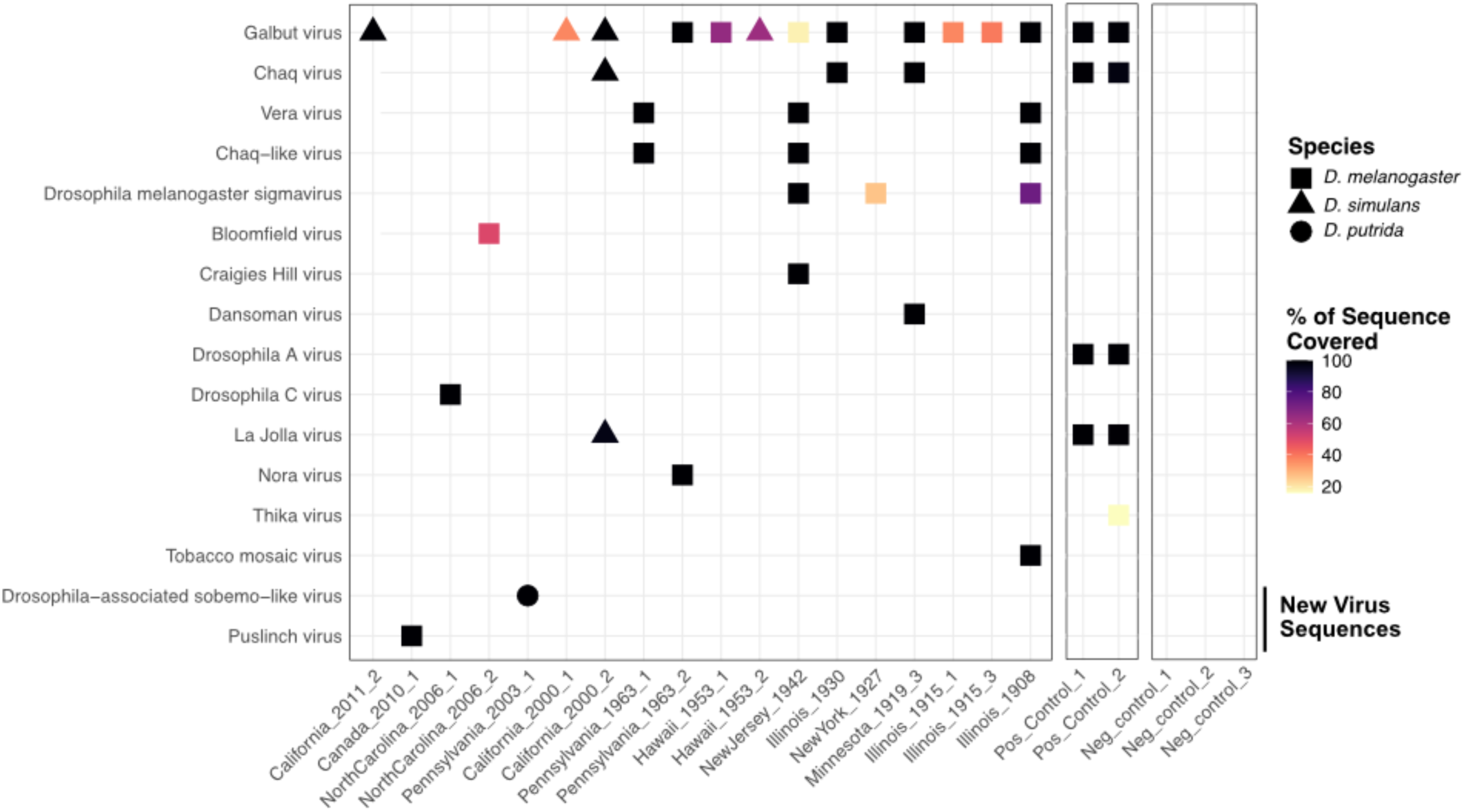
Entomological museum specimens harbored diverse viral sequences. Heatmap showing virus sequences recovered from individual museum samples. Shape corresponds to host species and color corresponds to the percent of reference sequence covered (**Table 2**). For segmented viruses, the average coverage across all segments is shown. Viruses detected in positive control datasets, made from pooled FoCo17 flies, are shown. No reads mapping to any of these viruses were present in any of the three negative control datasets.

We identified two coding-complete genomes from previously undescribed viruses. The first was a bunyavirus from a *D. melanogaster* collected in Ontario, Canada in 2010, which we named Puslinch virus. The Puslinch virus L protein shares only ∼35% pairwise amino acid identity with its nearest known relatives, which were viruses in the genus Herbevirus (**Fig. 4**; **Fig. 5**; **Table 2**). The other new virus sequence was a sobemo-like virus from a *D. putrida* collected in Pennsylvania, USA in 2003 (**Fig. 4**; **Fig 5**; **Table 2**).

**Figure 5:**
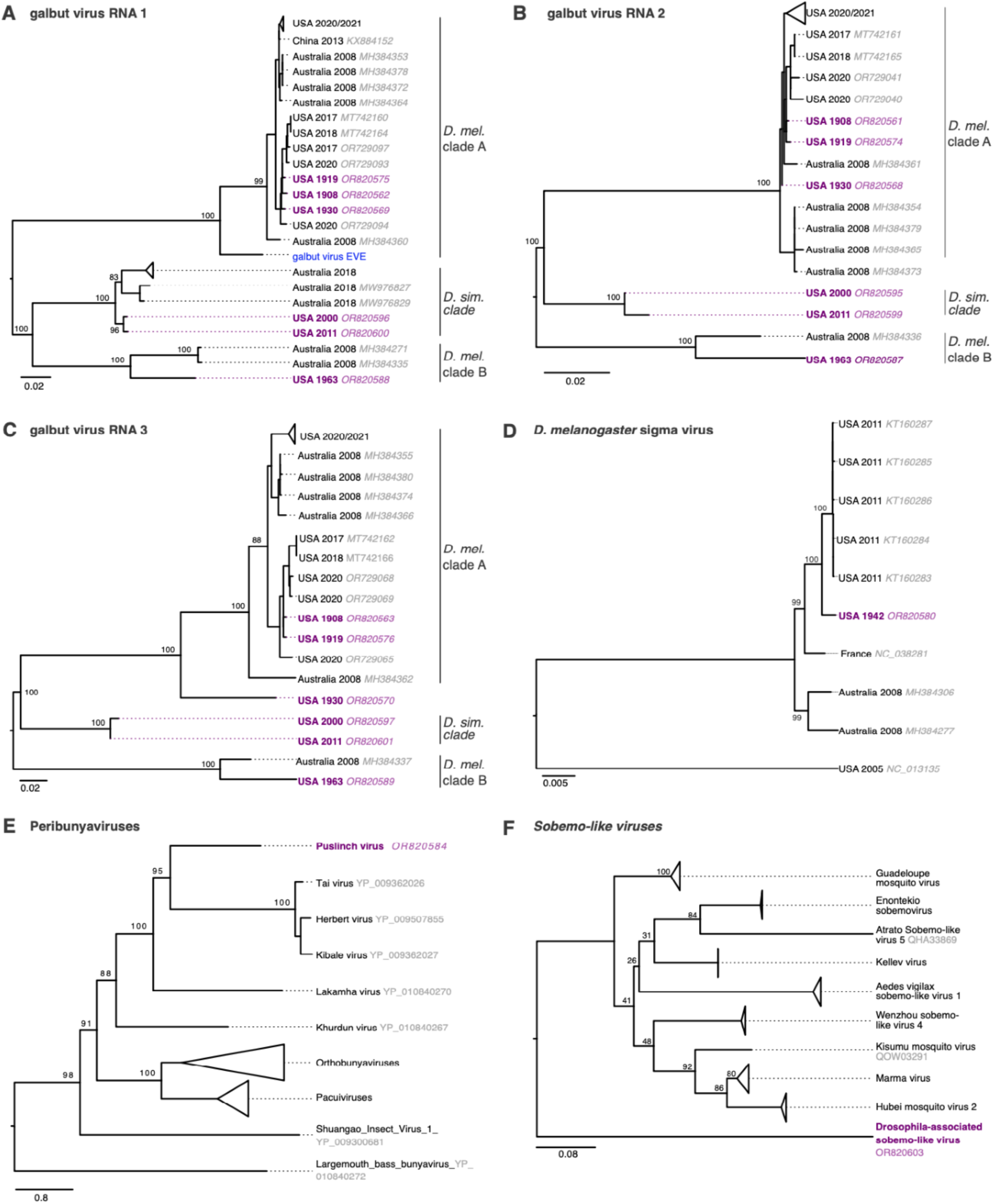
Trees showing museum sample virus sequences in the context of related contemporary sequences. (A) Maximum likelihood trees for galbut virus RNA 1 (RdRp) sequences, (B) galbut virus RNA 2 sequences, (C) galbut virus RNA3 sequences, (D) *D. melanogaster* sigma virus sequences, (E) L protein sequences for viruses in the family *Peribunyaviridae*, and (F) putative RNA-dependent RNA polymerase protein sequences related to the new *Drosophila*-associated sobemo-like virus. Trees in (A)-(D) are based on nucleotide alignments and trees in (E) and (F) are based on protein sequence alignments. Purple color indicates museum sample-derived sequence generated in this study. Accession numbers are noted except when groups of closely related sequences are collapsed. All trees are midpoint rooted. Blue tip indicates the sequence of an endogenized galbut virus RNA 1 sequence.

Additionally, we recovered a complete tobacco mosaic virus (TMV) genome with high coverage depth (850x) from the 1908 *D. melanogaster* specimen. We speculated that this sequence may have originated from contamination of the specimen, perhaps during handling by a smoker^41^.

Positive control libraries, constructed from RNA from pooled FoCo-17 flies, contained the expected virus sequences (**Fig. 4**). To avoid potential read contamination via index hopping, we sequenced positive control libraries on a separate sequencing run from the museum specimens^42^. Negative control datasets were included in each run and contained fewer than 15k reads after trimming of adapter and low-quality bases. Negative control datasets contained zero reads mapping to any of the virus sequences present in any of the other datasets (**Fig. 4**)^39^.

### Relatedness of old virus sequences to contemporary ones

Sequences from museum samples of previously described viruses ranged from 83.5% to 100% identical to existing sequences with 38% of the sequences sharing ≥ 99% pairwise identity to existing sequences (**Table 2**). Notably, all vera virus RNA 1 sequences, including one from 1908, were 100% identical to their closest available existing sequence, though the vera virus RNA 2 and chaq-like sequences from the same samples were not identical to existing sequences.

We initially considered a Bayesian approach to generate evolutionary rate estimates but found our data had insufficient temporal signal^43^. We instead generated initial estimates of evolutionary rates using differences in sequence identity and sampling times for the most closely related contemporary sequences (**Supp.** Fig. 7; **Table 2**,). This may underestimate rates due to saturation and might overestimate rates because the old sequences are likely not the direct ancestors of the contemporary sequences^44–46^. Sequences without a suitably similar contemporary sequence (e.g., USA 2000 and 2011 galbut virus *D. simulans* RNA 2 and 3) were not analyzed in this way. Estimated rates were generally higher in sequences from more recent samples (2000-2011) and lower in sequences from older samples, with sequences from 1908 showing the lowest rates (**Supp.** Fig. 7). This observation is consistent with the time-dependent rate phenomenon, in which evolutionary rate estimates decrease as the timescale of measurement increases^44,47^.

The most common contemporary viral infection of *D. melanogaster* is galbut virus, the success of which is likely attributable to efficient biparental vertical transmission and minimal apparent fitness costs^26,30,48^. Detection of galbut virus in museum samples by sequencing matched detection by RT-qPCR (**Table 1**). In maximum likelihood trees, three clades of galbut virus were evident: two clades (A and B) consisted of sequences from *D. melanogaster* while the third clade consisted of sequences from *D. simulans* (**Fig. 5A-C**). An endogenized galbut virus RNA 1 sequence was on its own branch most closely related to clade A (**Fig. 5A**)^49^. All the historic galbut virus sequences fell within existing galbut virus diversity and clustered with other sequences from the same host i.e., the two *D. simulans* sequences clustered with previously described *D. simulans* sequences and the *D. melanogaster* sequences with known *D. melanogaster* RNA1 sequences (**Fig. 5A**). Interestingly, the RNA1 and RNA2 sequence from Illinois 1930 clustered with clade A but this virus’s RNA 3 was situated on a separate branch. The 1930 Illinois RNAs 1 and 2 were 99% identical to contemporary sequences in GenBank while RNA 3 was only 89% identical. This phylogenetic discordance is consistent with galbut virus reassortment^50^.

The sigma virus sequence from 1942 clustered within the previously described diversity of *D. melanogaster* sigma viruses (**Fig. 5D**). Puslinch virus and *Drosophila*-associated sobemo-like virus sequences were situated on their own branches on trees of related sequences (**Fig. 5E-F**). These new virus sequences are highly divergent and may establish new genera or higher order taxa.

### Certain types of RNA survived better in old samples

We performed a variety of analyses to characterize the molecular and genomic properties of surviving RNA. An initial indication that not all types of RNA survived equally came from the strandedness of virus-mapping reads. We constructed libraries using a protocol that preserved information about the RNA strand from which reads derived^51^. Galbut virus is a partitivirus, which have dsRNA genomes^52^. But in infected flies, nearly all galbut virus RNA was positive (+) sense (98.2% in fresh frozen flies; **Fig. 6; Supp.** Fig. 8). This is presumably because most galbut virus RNA in infected cells is messenger RNA (mRNA). The fraction of +strand galbut virus RNA decreased in older samples. In experimentally dried samples an average of 89.1% of reads were from +strand RNA, and this value fell to 74.1% in museum specimens (p = 1.4x10^-4^; **Fig. 6**). The decreasing fraction of +strand galbut virus RNA could be explained by the preferential survival of double-stranded galbut virus RNA in older samples.

**Figure 6:**
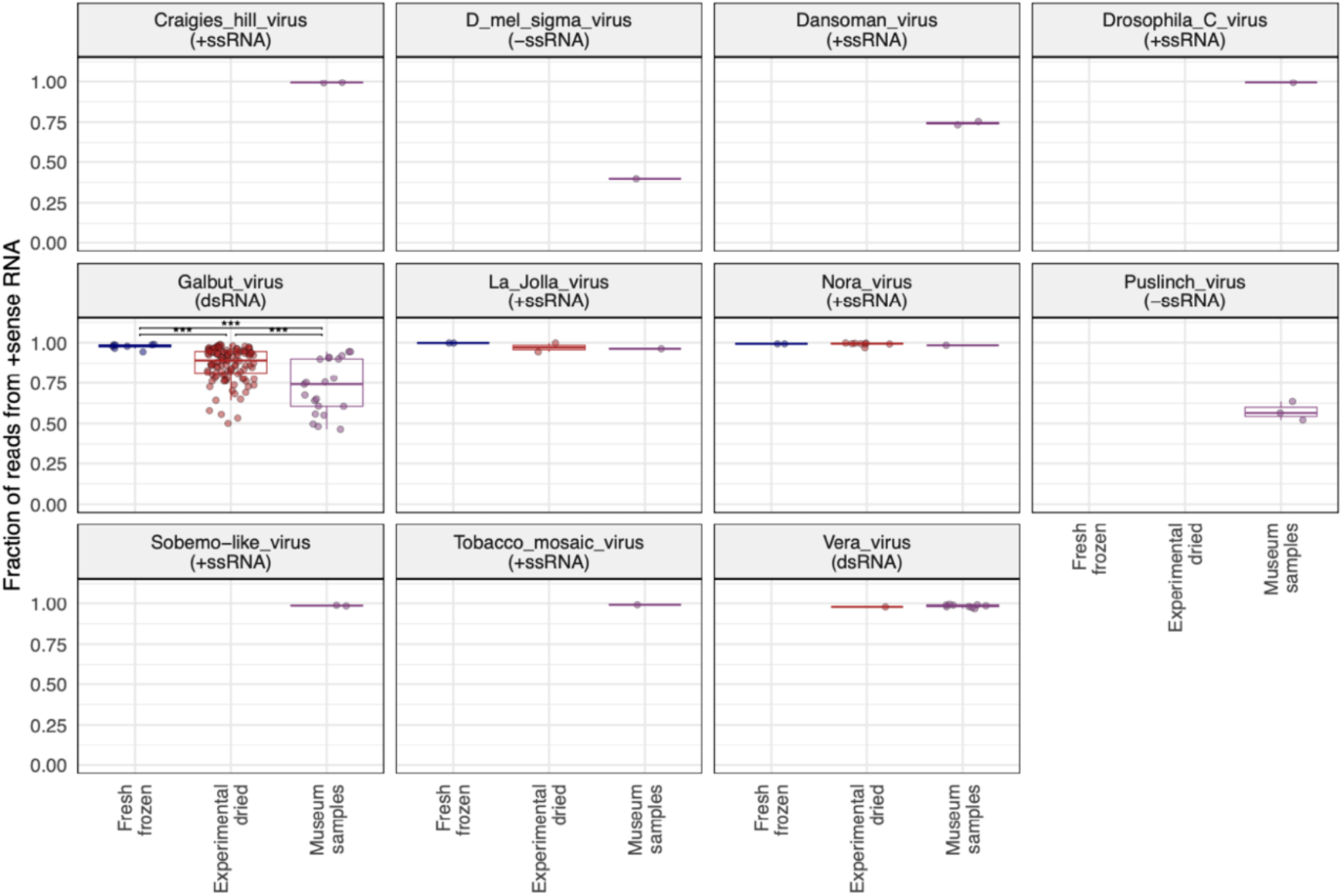
The strandedness of galbut virus RNA in old samples is consistent with preferential survival of dsRNA. The fraction of virus-mapping reads that originated from +strand RNA is plotted. Each point represents a dataset from an individual *D. melanogaster* fly. Significance levels of Wilcoxon test adjusted *p*-values are indicated: ns: p > 0.05; *: p <= 0.05; **: p <= 0.01; ***: p <= 0.001; ****: p <= 0.0001. Vera virus and other viruses were not present in sufficient numbers in different datasets to statistically evaluate how their strand ratios changed over time.

Coverage of galbut virus in museum datasets derived from all segments and was not concentrated in regions of RNA 1 corresponding to the PCR primers we use to detect galbut virus in our lab (**Supp.** Fig. 8). The lack of higher coverage in PCR product regions supports the conclusion that galbut virus sequences did not result from contamination by contemporary PCR products.

Other viruses had different fractions of +strand RNA that were consistent with the genome type of each virus (**Fig. 6**). For instance, a majority of reads from *D. melanogaster* sigma virus, a negative (-) strand RNA virus, were from -strand RNA^29^. RNA from Nora virus (a +strand virus) remained almost totally +strand in all samples^33^. The fraction of +strand vera virus RNA, another partitivirus, remained high in old samples (**Fig. 6**). This indicated that it was not simply dsRNA that survived in older samples.

Nevertheless, there was additional evidence from host-mapping reads that double-strandedness contributed to preferential RNA survival. Most host-mapping RNA derived from ribosomal RNA (rRNA), and most rRNA-mapping reads derived from the +strand (**Fig. 7**). In fresh samples, there was on average 698x more +strand rRNA than -strand rRNA. The small amount of -sense rRNA could have derived from anti-sense transcription of rRNA genes or pseudogenes^53^. In experimentally dried and museum samples, the +strand:-strand rRNA ratio dropped to 142:1 and 30:1, respectively (p = 8.2x10^-13^ and 2.8x10^-22^; **Fig. 7**). This drop was driven by a relative decrease in +strand rRNA and a relative increase in -strand rRNA in older samples (**Supp**. Fig. 9). This pattern could be explained by the preferential survival of sense:antisense rRNA duplexes. However, even in old samples, there was 30x more +strand rRNA than –strand (**Fig. 7**), so sense:antisense duplexes could not account for all surviving rRNA.

**Figure 7:**
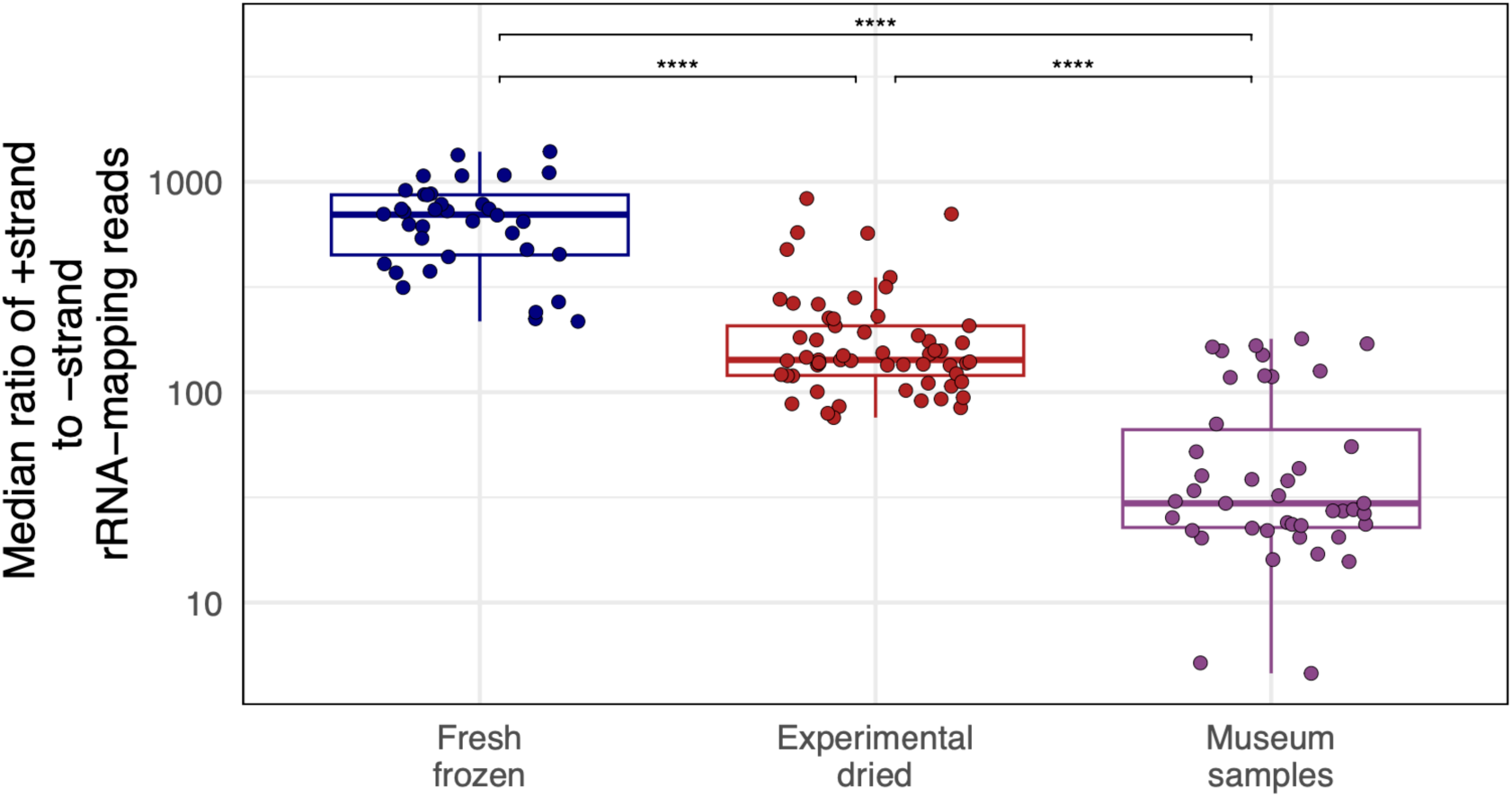
Strand ratios of surviving ribosomal RNA were consistent with preferential survival of dsRNA. The median ratio of +strand to –strand coverage of rRNA-mapping reads. Points represent individual-fly datasets. Adjusted *p*-values significance levels from Wilcoxon test are indicated.

Coverage levels across rRNAs varied more in old samples than they did in new samples, where coverage was relatively even (**Fig. 8**). For instance, coverage surrounding position 1500 of the 18S rRNA and position 3000 of the 28S rRNA had consistently lower coverage in old samples (**Fig. 8**). In contrast, in fresh samples coverage over these same regions was similar to average coverage.

**Figure 8:**
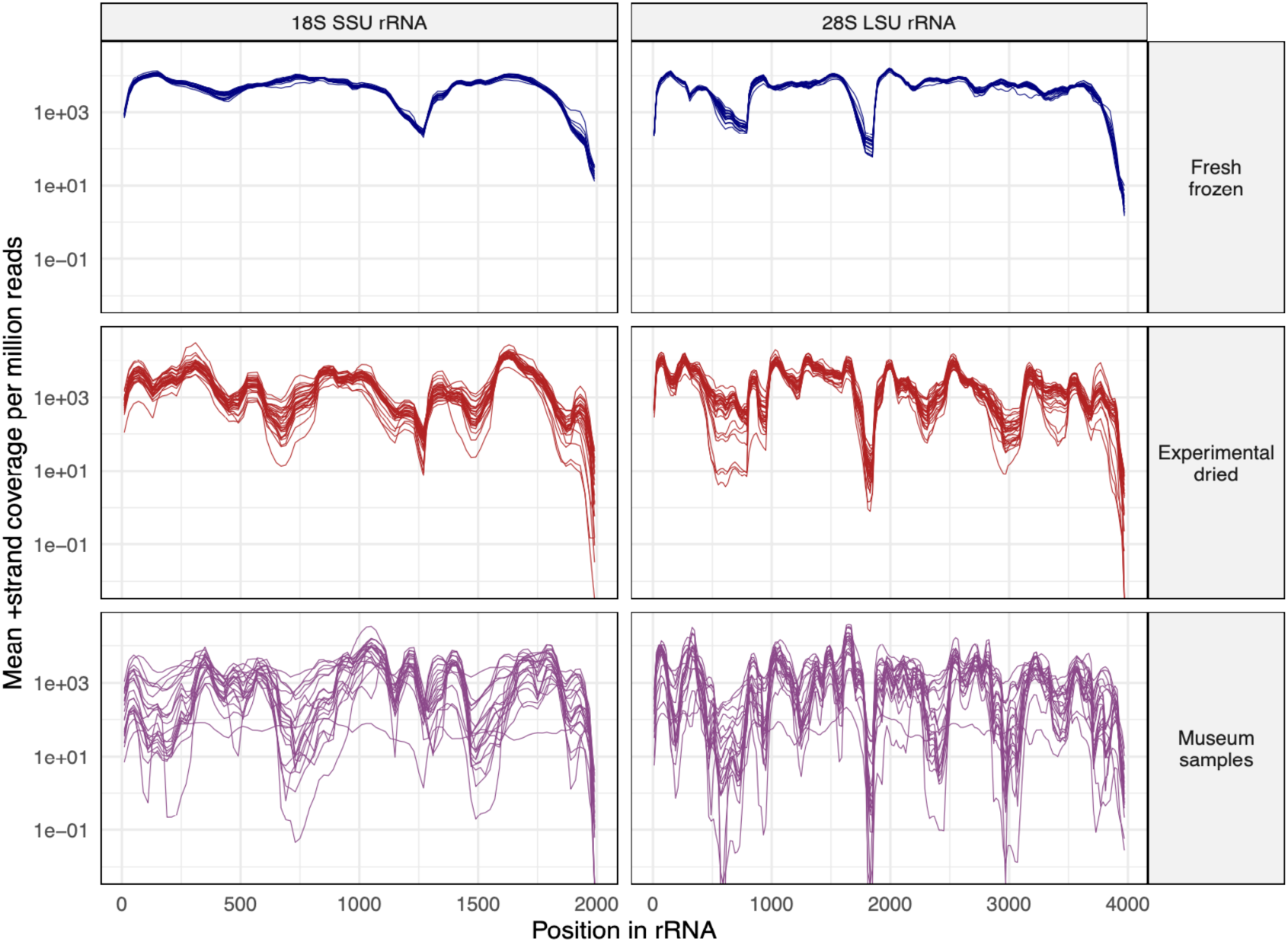
Certain regions of ribosomal RNA were underrepresented in old samples. The mean +strand rRNA coverage per million reads in 20 bp windows is plotted for *D. melanogaster* datasets. Each line represents a dataset from an individual fly. SSU: small subunit (18S) rRNA; LSU: large subunit (28S) rRNA.

To investigate why some rRNA regions might not survive as well as others, we took advantage of a high resolution cryo-electron-microscopy structure of the *D. melanogaster* 80S ribosome^54^. This structure includes ribosomal proteins and RNA (**Fig. 9A**). It is possible to resolve individual rRNA bases in the structure and to determine whether they are interacting with other bases, and whether they are present in the structure (**Fig. 9B**). We used this structure and RNApdbee software to individually assign the 5965 rRNA bases to one of four categories: paired with other rRNA bases (62.9%), unpaired (29.2%), present in higher-order secondary structures like pseudoknots (4.5%), or missing from the 3D structure altogether (3.4%; **Fig. 9B**)^54–56^.

**Figure 9:**
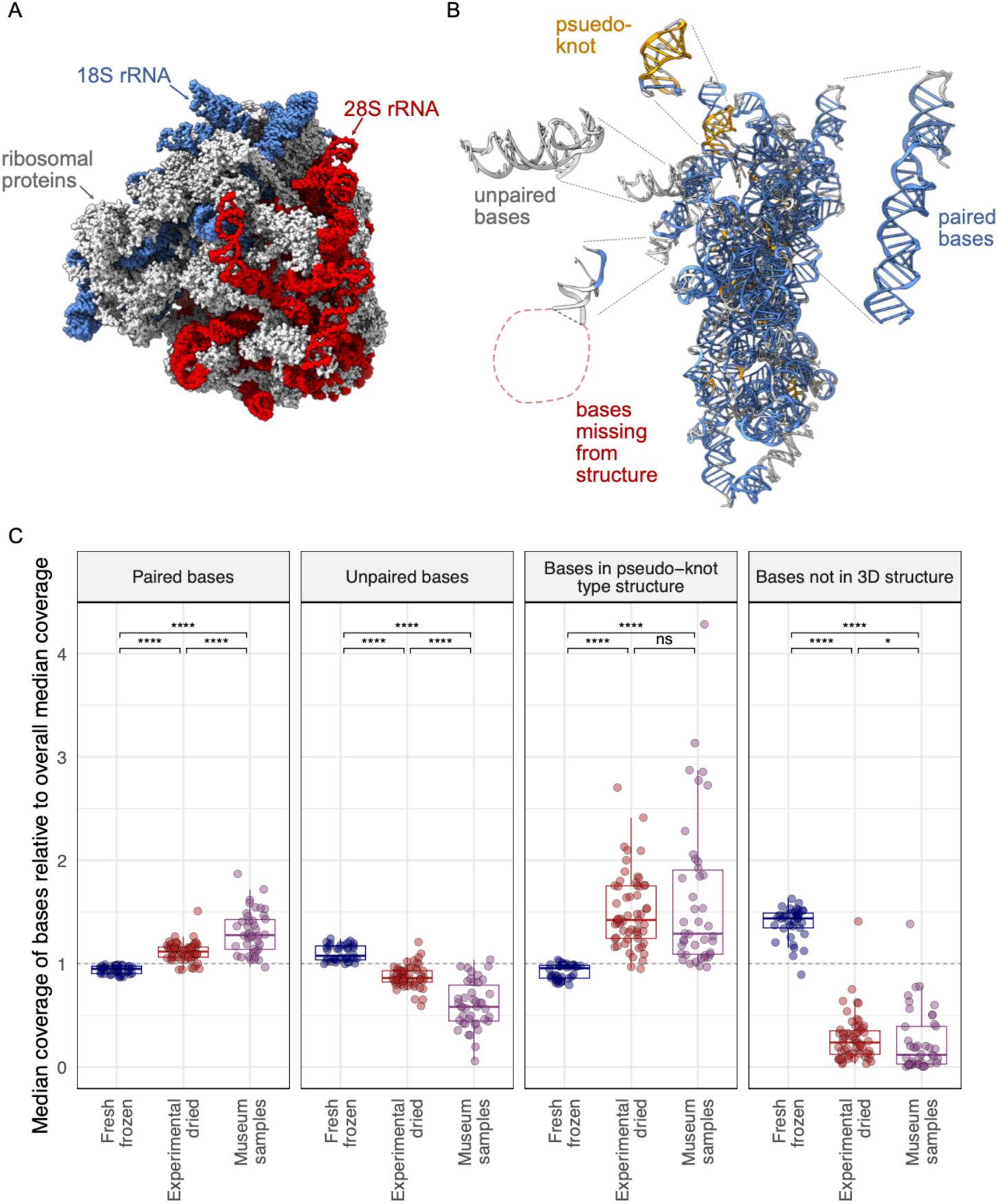
Different types of ribosomal RNA exhibited differential survival in old samples. (A) The structure of the *D. melanogaster* 80S ribosome from Anger at al, showing ribosomal proteins and the 18S and 28S ribosomal RNAs. (B) The 18S rRNA with individual bases color coded to indicate whether they are paired with other rRNA bases (blue), unpaired (grey), or involved in higher order pseudo-knot type structures (gold). Some bases were not captured in the structure (red). Bases were binned into categories using RNApdbee software. The 18S rRNA is rotated relative to panel A. (C) The median coverage level of bases in the indicated categories relative to total median coverage is plotted. Each point represents a dataset from an individual *D. melanogaster* fly. Significance levels of Wilcoxon test adjusted *p*-values are indicated.

We calculated coverage of bases in each of these four categories relative to total median coverage (**Fig. 9C**). Paired bases had higher average coverage in older samples (p = 1.9x10^-19^ in museum samples vs. fresh ones). Similarly, bases in higher-order secondary structures like pseudoknots had higher coverage in old samples (p = 3.3x10^-17^ relative to fresh). In contrast, unpaired bases had lower average coverage in old samples (p = 3.8x10^-20^ vs. fresh samples). Bases that were not captured in the 3D structure were the least well represented in older samples, with coverage levels 8x lower than overall average coverage in museum samples (p = 1.0x10^-19^ relative to fresh frozen). Experimentally dried sample coverage levels were generally intermediate between fresh and museum samples, consistent with their intermediate age (**Fig. 9C**).

These patterns suggested that the molecular environment surrounding rRNA bases influenced the likelihood that they would survive. Being base-paired or in a higher-order RNA secondary structure protected bases, enabling them to survive over longer periods. In contrast, unpaired bases or bases not present in the 3D structure - presumably because they were outside of the protective environment of the ribosome - were less likely to survive.

### Different types of non-ribosomal RNA survived differentially in old samples

We also quantified the extent to which different types of non-ribosomal host RNA survived in old samples. We mapped reads from *D. melanogaster* samples to the *D. melanogaster* genome and used the ALFA software to quantify the different types of host RNA present in each dataset^57^. ALFA combines mapping information with genome annotation to assign mapped reads to one of a dozen RNA types (**Fig. 10**). As expected, most reads were categorized as rRNA, though there was relatively less rRNA in older samples (**Fig. 10A**; p = 5.1x10^-6^ for museum vs. fresh samples). There was also relatively less protein-coding mRNA in older samples (**Fig. 10B**; p= 5.7x10^-4^). Opposite strand RNA levels – that is, reads derived from the RNA strand opposite an annotated feature - were elevated in older samples, consistent with the preferential survival of sense:antisense transcript duplexes (p = 1.8x10^-7^). Highly structured transfer RNA (tRNA) levels were also elevated in older samples (p = 5.9x10^-5^)^58,59^.

**Figure 10:**
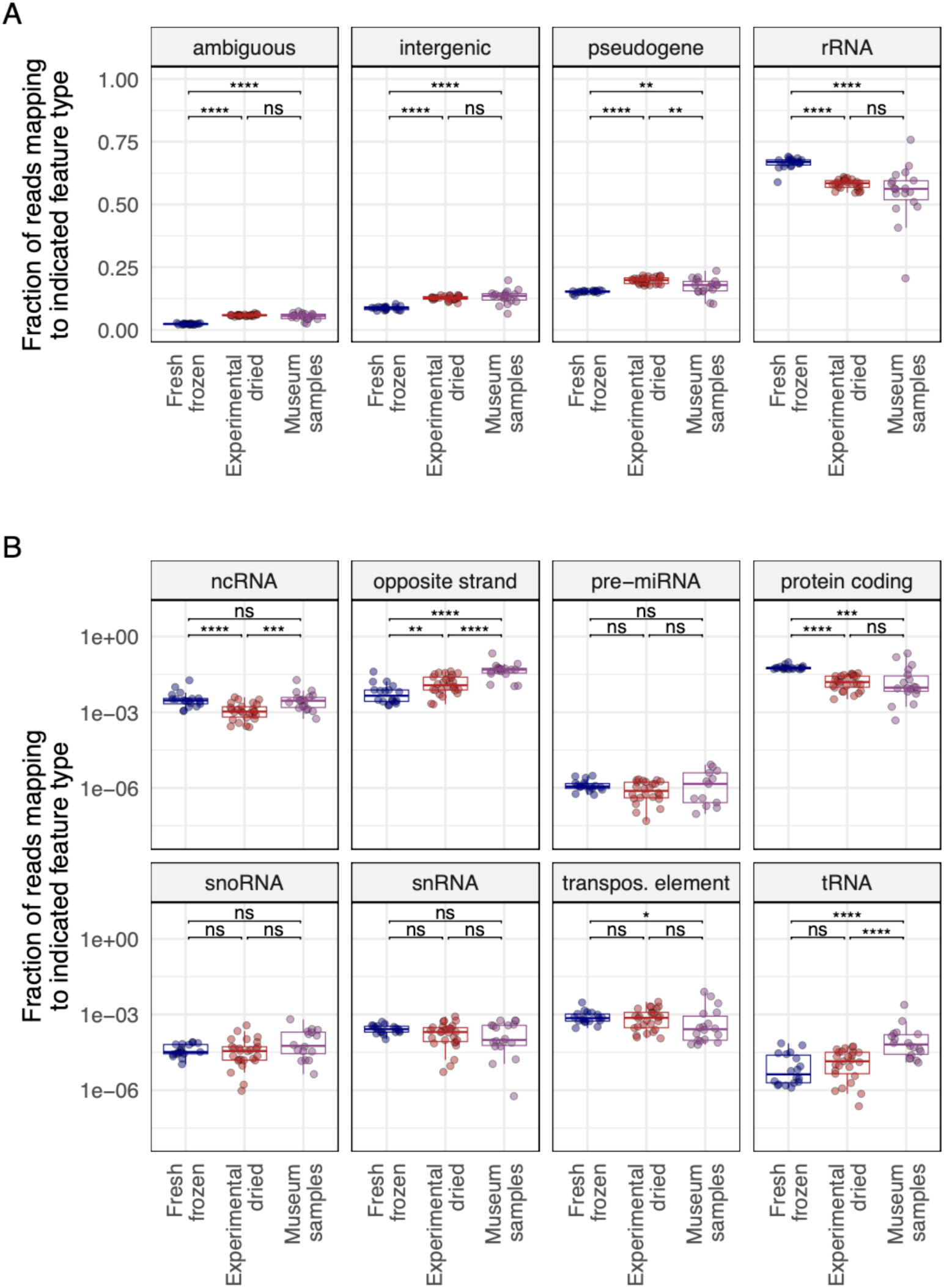
Certain types of RNA survived better in old samples. The fraction of reads in *D. melanogaster* datasets mapping to the indicated RNA types is plotted. Each point represents a dataset from an individual fly. Adjusted *p*-values significance levels from Wilcoxon test are indicated. (A) Types of RNA present at >5% median abundance. (B) Types of RNA present at <5% abundance. Abbreviations for different RNA types: ncRNA: non-coding RNA; miRNA: micro RNA; snoRNA: small nucleolar RNAs; snRNA: small nuclear RNA; tRNA: transfer RNA.

### Old RNA was chemically damaged

Old DNA molecules are fragmented and chemically damaged^60^. Polymerases can misincorporate bases when copying damaged templates, causing damage to manifest as substitutions in library molecules^61–63^. For example, deaminated cytosines, common in ancient DNA, result in C-to-T substitutions in library molecules. In fact, signatures of chemical damage in sequence reads are taken as evidence that reads derive from old DNA and not from contaminating contemporary DNA.

We quantified mismatch patterns in mapped reads to look for signatures of chemical damage in old RNA (**Fig. 11**). Mismatch frequencies varied in different sample types. Some differences in mismatch frequencies could be attributed to samples being sequenced on different sequencing runs (we sequenced museum samples separately from other samples to avoid the possibility of read misassignment from index-hopping) ^64^. The largest difference between old and new samples was an elevated frequency of C-to-T substitutions in reads from museum samples. The median level of C-to-T mismatches in datasets from museum samples was 40x higher than in fresh-frozen datasets (2.0x10^-3^ frequency of C-to-T mismatches vs. 5.0x10^-5^ (**Supp.** Fig. 10; p=1.0x10^-10^). This increased frequency of C-to-T substitutions is consistent with deamination of cytosines in old RNA.

**Figure 11.**
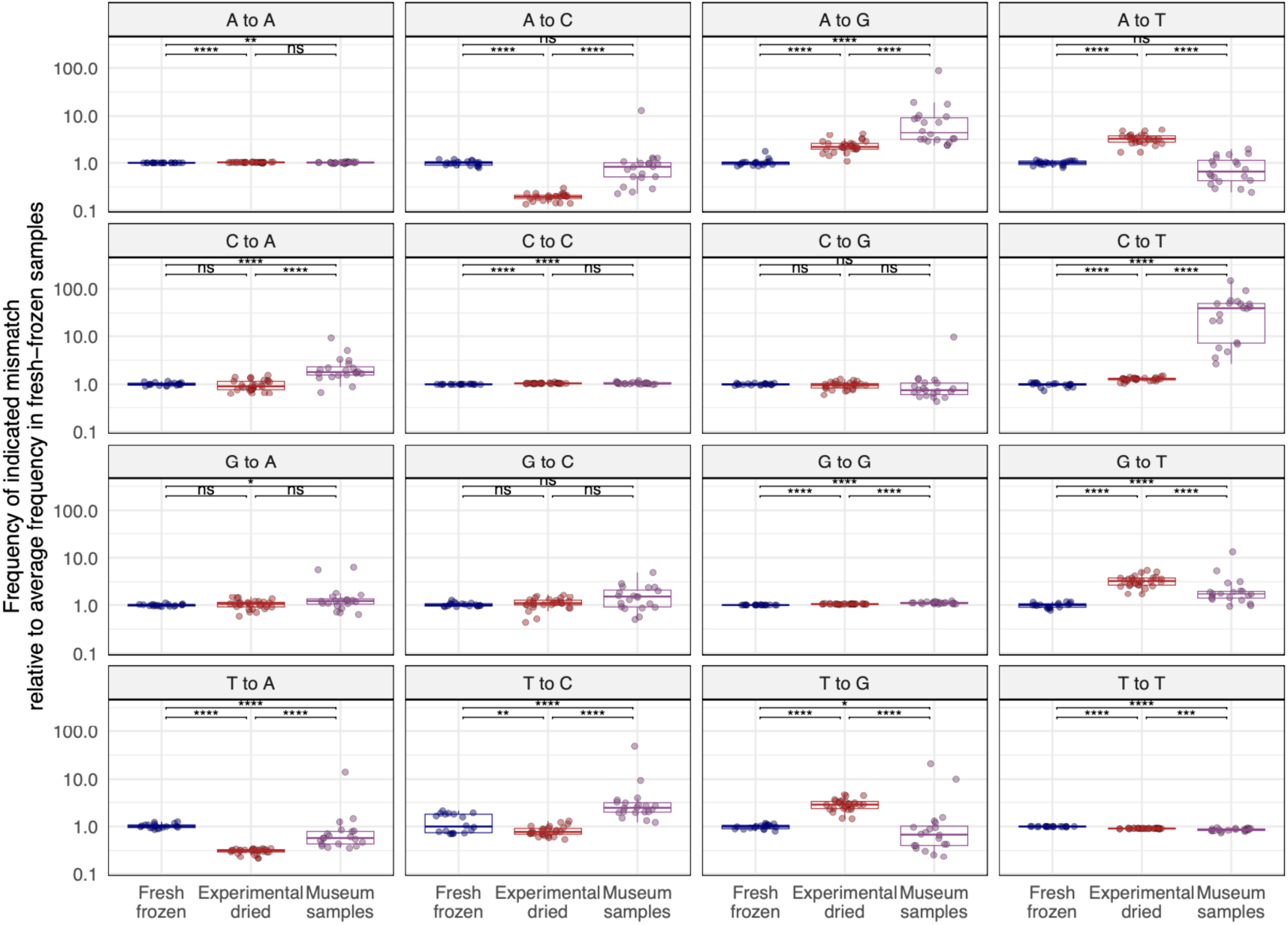
Old RNA was chemically damaged and exhibits mismatch patterns consistent with cytosine and adenine deamination. Mismatches in rRNA-mapping reads from *D. melanogaster* datasets were quantified and the frequency of each mismatch type relative to the median frequency in fresh-frozen datasets is plotted. Each point represents a dataset from an individual fly. Adjusted *p*-values significance levels from Wilcoxon test are indicated.

A-to-G substitutions were the next most elevated type of mismatch in old samples (**Fig. 11**). A-to-G substitutions occurred at a rate 4.4-fold higher in museum samples than in fresh datasets (5.7x10^-4^ vs. 1.3x10^-4^; p=1.6x10^-10^). Such substitutions can result from spontaneous deamination of adenine to hypoxanthine^60,65^.

### Host-mapping reads exhibited species specificity

We took advantage of the fact that our musuem samples derived from 4 different species to investigate the possibility that host-mapping reads – specifically rRNA-mapping reads – might result from contemporary contamination. The likely source of fly-mapping contamination would be the *D. melanogaster* that we rear and study in our lab^48^. The 18S and 28S ribosomal RNA sequences of *D. melanogaster* and *D. simulans* share over 99% pairwise identity which makes them difficult to distinguish by competitive read mapping. We therefore collected and mapped reads to the relatively variable region of the ribosomal RNA between the 18S and 5.8S genes (the internal transcribed spacer 1, ITS) of the 4 fly species that we sampled (**Table S1**; **Fig. 12; Supp.** Fig. 11). All ITS-mapping reads in museum datasets mapped to the expected species ITS sequence and no reads mapped to an unexpected species (**Fig. 12**). Positive and negative control samples behaved as expected, with water negative control datasets containing no reads mapping to any of these 4 ITS sequences (**Supp.** Fig. 11). If fly rRNA-mapping reads were from contamination, we would have expected there to be *D. melanogaster*-mapping reads in the non-*melanogaster* and negative control datasets; instead, there were none. The perfect concordance of ITS mapping and sample species supported the conclusion that the host-mapping reads in our datasets were indeed from museum samples.

**Figure 12:**
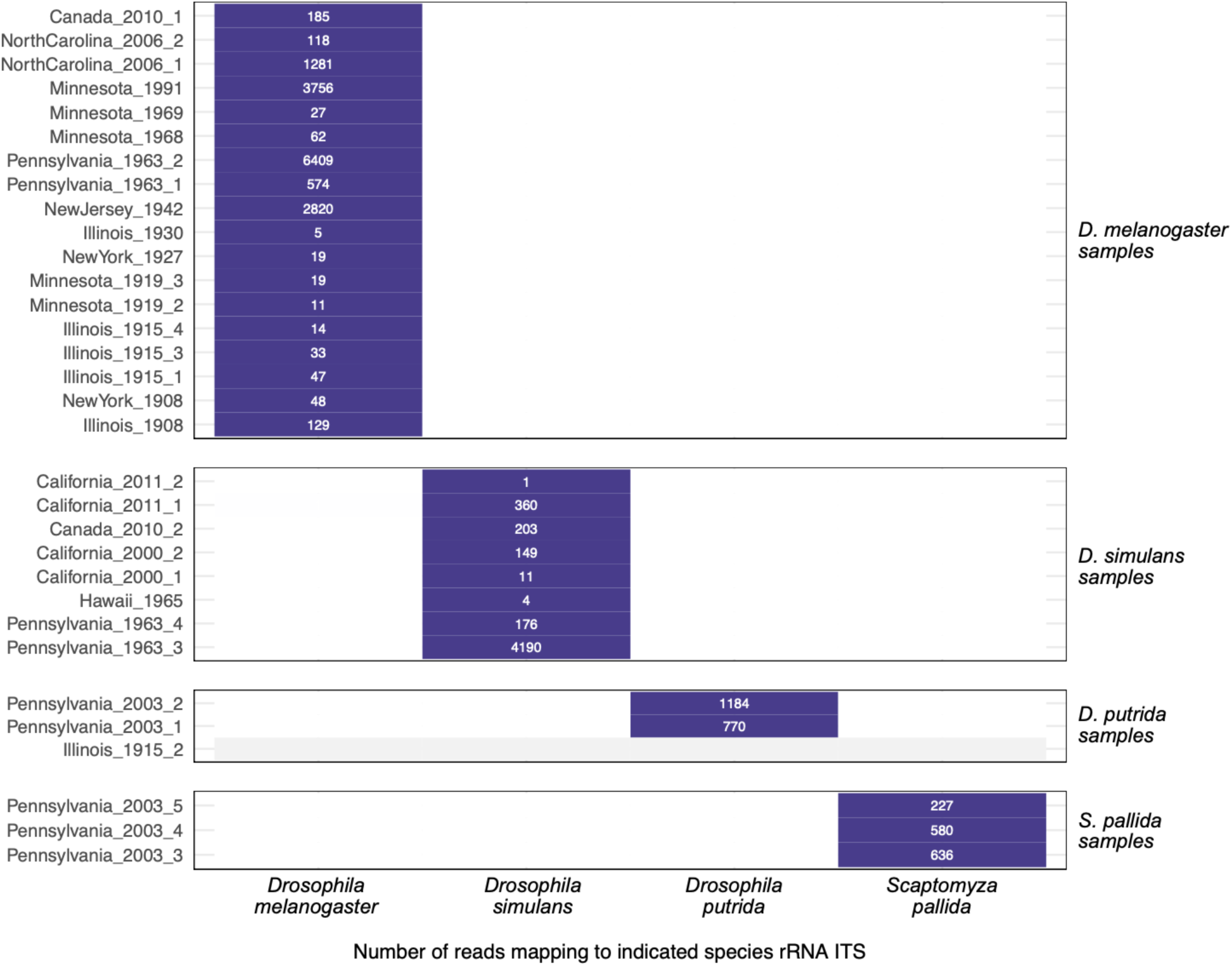
Host-mapping reads exhibited species specificity. The number of reads mapping to the indicated species rRNA internal transcribed spacer 1 (ITS) sequence for each museum sample dataset is shown. Samples are grouped according to their molecular species assignment (**Table S1**). Datasets with no ITS-mapping reads are represented with grey.

## DISCUSSION

In this study we took a two-pronged approach to investigate RNA stability in old biological samples. In flies and mosquitoes stored dry at room temperature for weeks or months, RNA grew increasingly fragmented, but RNA yields did not significantly decrease, and cellular and viral RNA remained detectable. Perhaps the most unexpected result from this experiment was the limited difference in RNA quality and quantity between dried and frozen samples (**Fig. 1; Supp.** Figs. 2**, 4**). We recovered RNA from dried flies that were from as far back as the 1890s and used this RNA to make sequencing libraries. We recovered coding complete virus sequences from 11 old specimens and identified two novel virus sequences. We also showed that certain types of RNA tended to survive in old samples, including dsRNA and RNA in secondary structures or ribonucleoprotein complexes.

There were limitations to the experimental arm of our study. First, RNA recovery from individual flies and mosquitoes - even fresh ones - was variable. Second, we used outbred populations of flies and mosquitoes with different levels of virus infection between individuals. In future studies, optimization of our RNA isolation protocol to decrease yield variability between individual flies and mosquitoes would be useful. It would also be better to use pinned specimens from inbred populations with less variable infection phenotypes.

We found that not all RNA survived equally well and described molecular features that influenced how well different types of RNA survived. dsRNA survived better than ssRNA. Surviving RNA duplexes derived from intermolecular base pairing, such as sense-antisense duplexes (**Figs. 6, 7A, 10B**) and from intramolecular base pairing, such as paired rRNA bases (**Fig. 9C**). The preferential survival of dsRNA likely reflects decreased susceptibility to hydrolysis^66^. However, double strandedness could not explain all RNA survival, as RNA from certain viruses remained largely +strand (**Fig. 6**) and unpaired rRNA survived (albeit less than paired rRNA; **Figs. 7, 9**). It may be that ribonucleoprotein complexes like virus particles and ribosomes provide a protective environment in which RNA can survive longer.

Like old DNA, old RNA was fragmented and chemically damaged (**Figs. 2C, 11**). And, like DNA, deamination was the most evident type of damage (**Fig. 11**). Other types of damage may be common in old RNA but undetectable by sequencing. The extent to which ribonucleases, deaminases, and other enzymes involved in normal RNA processing continue to function in dead cells is unclear, but it is likely that most RNA fragmentation and damage in old samples results from spontaneous chemical rather than enzymatic degradation.

Environmental factors like temperature and humidity likely play a major role in the survival of RNA. It was notable that none of the 12 specimens from tropical Hawai’i yielded enough RNA to be detected fluorometrically and only two generated sequenceable libraries (**Fig. 2b, Table 1 & Supp.** Fig. 5). Humidity might be more important than temperature, as RNA was recovered from 1000-year-old corn in Arizona, where temperatures fluctuate to extremes, but humidity is very low^15^. If this were true, RNA might survive better in drier environments and in quickly desiccating samples like the small insects that are the focus of this work. Even for much larger samples like wooly mammoths, dehydration has been proposed to promote preservation of nuclear architecture and biological macromolecules^67^.

The virus sequences we recovered revealed that the *D. melanogaster* virome has not changed much over the last 120 years. All the viruses that we detected in *D. melanogaster*, except for Puslinch virus, had already been described in contemporary *D. melanogaster*^25,26^. For the most part the old sequences were closely related to contemporary sequences (**Table 2**). For example, the viruses infecting the fly collected in 1908 were >98% identical to known *D. melanogaster* viruses (**Fig. 4**; **Table 2**).

The discrepant phylogenetic positions of the 1930 galbut virus segments is consistent with reassortment^68^. The 1930 galbut virus RNA 3 sequence that is positioned on its own branch may correspond to an extinct RNA 3 lineage, or simply one that has not been sampled before (**Fig. 5C**). Similarly, the endogenous galbut virus RNA 1 sequence in these trees may represent an extinct lineage or an ancestor of contemporary galbut virus RNA 1 sequences (**Fig. 5A**)^49^. A potential benefit of large-scale sequencing of old samples is that it could provide estimates of how frequently virus lineages go extinct from a particular host.

Contamination from contemporary nucleic acid has been a long-standing concern for those sequencing ancient DNA^69^. There was evidence of low-level contamination in our datasets (**Fig. 3**), and it is difficult to completely avoid contamination in NGS datasets, even when sequencing new samples^70,71^. But, there were numerous lines of evidence to support the idea that our conclusions were based on sequences from actual old RNA and not from contemporary contamination. Old RNA was short and chemically damaged; properties that are shared with old DNA^60^. Characteristics of RNA from experimentally dried samples were generally intermediate between those of fresh RNA and museum RNA, consistent with their intermediate age (**Figs. 6-11**). We recovered different virus sequences from different old samples, and reads in individual datasets supported the same distinct virus sequences (**Table 2**). The virus sequences we recovered were for the most part known *Drosophila*-infecting viruses. Although many virus sequences from old samples were similar to existing contemporary sequences, most were not identical (**Table 2**). The exceptions were three vera virus RNA 1 sequences that were identical to existing sequences. But the RNA 2 and chaq-like sequences from these samples were not identical to any contemporary sequences. The two new virus sequences we recovered were divergent, but their closest relatives were from other insects (**Table 2**; **Fig. 5**). The tobacco mosaic virus sequence from the 1908 dataset was an exception, which we can’t explain except to suggest that this specimen was contaminated during handling by a smoker, as individual cigarettes were found to contain as many as 10^9^

TMV RNA copies^41^. There was no peak in galbut virus coverage corresponding to our diagnostic RT-qPCR amplicons, which would be a prime candidate source of contaminating galbut virus sequences (**Supp.** Fig 6)^30,48^. All museum samples that were positive for galbut virus by RT-qPCR and generated a sequenceable library were also positive by sequencing (**Supp. Table 1**). Negative control datasets were uniformly free of virus and rRNA-mapping reads (**Fig. 4**; **Supp Fig. 11**). Galbut virus sequences from *D. simulans* museum samples clustered with contemporary *D. simulans* sequences, and the same for *D. melanogaster* sequences (**Fig. 5**).

Indeed, the species-specificity of host and virus sequences (**Figs. 5 & 12**) provides some of the strongest support for the idea that our analyses are based on nucleic acid from the actual old samples. If our conclusions were based on contaminating contemporary sequences, it would be necessary to invoke a complicated mechanism where contaminating host and virus nucleic acids sorted themselves by species into each sample (**Figs 5 & 12**). The overall apparent lack of contaminating virus and host sequences was consistent with the fact that all extractions were done in biosafety cabinets following a pre-PCR to post-PCR workflow, including DNase treatment before RT-qPCR and library preparation.

RNA can persist in biological samples over decades or centuries without freezing or fixation. The half-life of RNA in dead cells is clearly much longer than in living cells. The millions of dried specimens in museums represent a valuable source of RNA for reconstructing historic virus-host interactions^14,15^. It is tempting to speculate that the long-term survival of certain RNA might approach that of DNA^72^. It is time to move beyond the incorrect idea that old, dried samples are not a good source of useful RNA.

## MATERIALS AND METHODS

### Sample Collection

*Pinned Specimens:* FoCo-17 *D. melanogaster* were collected and pinned using standard entomological collection techniques with storage at room temperature or stored frozen at -80°C ^73^. Poza Rica *Aedes aegypti* mosquitoes were reared using standard lab techniques until adulthood^74^. Prior to blood feeding mosquitoes were collected and pinned or frozen.

*Museum Specimens:* Samples were obtained from museums and other institutions containing entomological specimens located in the United States of America and Canada. Altogether, 46 drosophilid samples, mainly *D. melanogaster* or *D. simulans*, were selected for the study. Museum species assignments were checked using molecular data as described below. One sample from California, California_2000_2, was initially identified as *D. melanogaster*, but our sequence data indicated that it was *D. simulans* (these species can be difficult to distinguish morphologically). Location, date of sample collection, sample storage type, extraction method, RNA quality and RNA extraction concentration are summarized in **Figure 2, Table 1** and **Supplemental Table 1**

### Sample Workflow

Deliberate protocols were used to minimize contamination during sample processing and library construction. Sample processing and library construction moved from dedicated pre-PCR rooms to a post-PCR room. RNA extractions were performed in a dedicated pre-PCR sample extraction room in a Class II type B2 biosafety cabinet which protects both personnel and samples. Initial library preparation steps, including reverse transcription, ligation, and setup of library amplification reactions were completed in a separate PCR setup room in an AirClean PCR workstation equipped with a HEPA filter (AirClean Systems, AC600 Series). Finally, library amplification and subsequent cleanup and pooling steps were performed in a third post-PCR lab space.

### RNA Extraction

*Non-Destructive Sampling:* For samples for which destructive sampling was prohibited, we adapted the protocol described by Santos et. al., 2018^35^. Briefly, individual samples were placed in a PCR tube containing 200µL of solution containing 200mM Tris HCl, 250mM NaCl, 25mM EDTA, 0.5% SDS and 400 µg/mL proteinase K. Samples were then incubated at 56°C for 16 hours. After incubation the supernatant was moved to a new tube for RNA isolation and the specimen was transferred to a cryovial containing 80% ethanol. RNA was purified using the Zymo RNA Clean & Concentrator Kit as described below.

*Ethanol Stored Sample Rehydration:* Four samples arrived in 70% ethanol. To prepare these samples for extraction, each specimen was rehydrated using a gradient of decreasing concentrations of ethanol. The specimens were suspended in 200µL 70%, 50%, 30% and 10% ethanol for 15 minutes on ice with agitation every 2-3 minutes prior to a final incubation in water for 15 minutes before RNA extraction following the destructive sampling method described below.

*Destructive Sampling:* A phenol/chloroform approach with mechanical disruption was used for RNA isolation of the experimentally dried and frozen samples as well as the museum specimens that could be destructively sampled. Briefly, each specimen was placed in a 2.0mL tube with a small BB (McMaster Carr, 1598K22) and 500µL of Trizol was added and incubated for 5 minutes. Tubes were then shaken in a TissueLyser (Qiagen, 9003240.) at 30 Hz for 3 minutes. 100µL of chloroform was added, thoroughly mixed and incubated for two minutes prior to being spun at 12,000xg for 10 minutes. The aqueous phase was retained for RNA purification using the Zymo RNA Clean & Concentrator-5 kit following manufacturer directions for the capture of small RNA fragments with slight modification (Zymo Research, R1013). Briefly, 225µL of sample was mixed with 225µL 100% ethanol and 225µL RNA binding buffer. Samples were transferred to spin columns with collection tubes and spun. To capture small RNAs, the flow through from the first spin was retained and mixed with an equal amount of 100% ethanol. This was transferred to the spin column and centrifuged. A wash using RNA wash buffer was done prior to a DNAse treatment following manufacturer recommendations (Zymo Research, E1010). After DNAse treatment, two more washes were complete prior to elution in 30µL of nuclease free water. All spins were performed at 9,000xg for 1 minute unless otherwise noted. For all extraction types, FoCo-17 male and female flies were used as a positive control and an empty tube was used as a negative control.

### RNA Quality and Quantification

RNA quality was assessed using a nanodrop spectrophotometer (Thermo Scientific ND2000USCAN) and RNA concentration was determined using the Qubit high-sensitivity RNA reagent (ThermoFisher, Q32852). RNA length was measured using the Agilent Tapestation High Sensitivity RNA screening tapes and reagents (Agilent, 5067-5579 & 5067-5580).

### RT-qPCR

*Pinned Samples:* Several targets were assessed for presence/absence and evidence of degradation using RT-qPCR (primer sequences in **Supplemental Table 3**). cDNA was generated using 5.5µL RNA, 10µM random hexamer primers (IDT), 10µM dNTPs and water to 13µL and incubated at 65°C for 5 minutes. Next, 4µL 5X FS buffer, 1µL 0.1M DTT and 1µL reverse transcriptase was added and incubated at 50°C for 60 minutes followed by a heat inactivation step at 80°C for 10 minutes after which each sample was diluted in 90µL nuclease-free water. qPCR was conducted using NEB Luna Universal qPCR Mastermix following manufacturer recommendations and cycling conditions (NEB, M3003).

### PCR controls

RNA from a pool of FoCo-17 flies was reverse transcribed to create a cDNA positive control for fly samples. For mosquito RT-qPCR, positive control cDNA was reverse transcribed from individual colony mosquitoes. We included these positive controls and three negative controls in all qPCR runs. Negative controls consisted of an extraction blank (a no sample extraction), a no-sample RT control, and a no template qPCR control. The use of negative controls at each step would have allowed us to determine when cross-contamination had been introduced if it had been. Positive/negative status was determined using the melt curve (within 1 degree of the positive control) for each sample and target. Agarose gel electrophoresis of qPCR products was used to determine positive/negative status when Ct and melt curve were ambiguous.

*Museum Collections:* We used RT-qPCR to determine if any samples had detectable levels of galbut virus RNA and RpL32 mRNA. To increase our ability to detect highly fragmented RNA, we designed a second set of primer that target a small region of the galbut virus RNA 1 and *D. melanogaster* mRNA RpL32. A 2% ethidium bromide agarose gel was used to confirm positive/negative status. Galbut virus positive female and male FoCo-17 flies were used as positive controls and water was used as a negative control.

**Supplemental Table 3:**
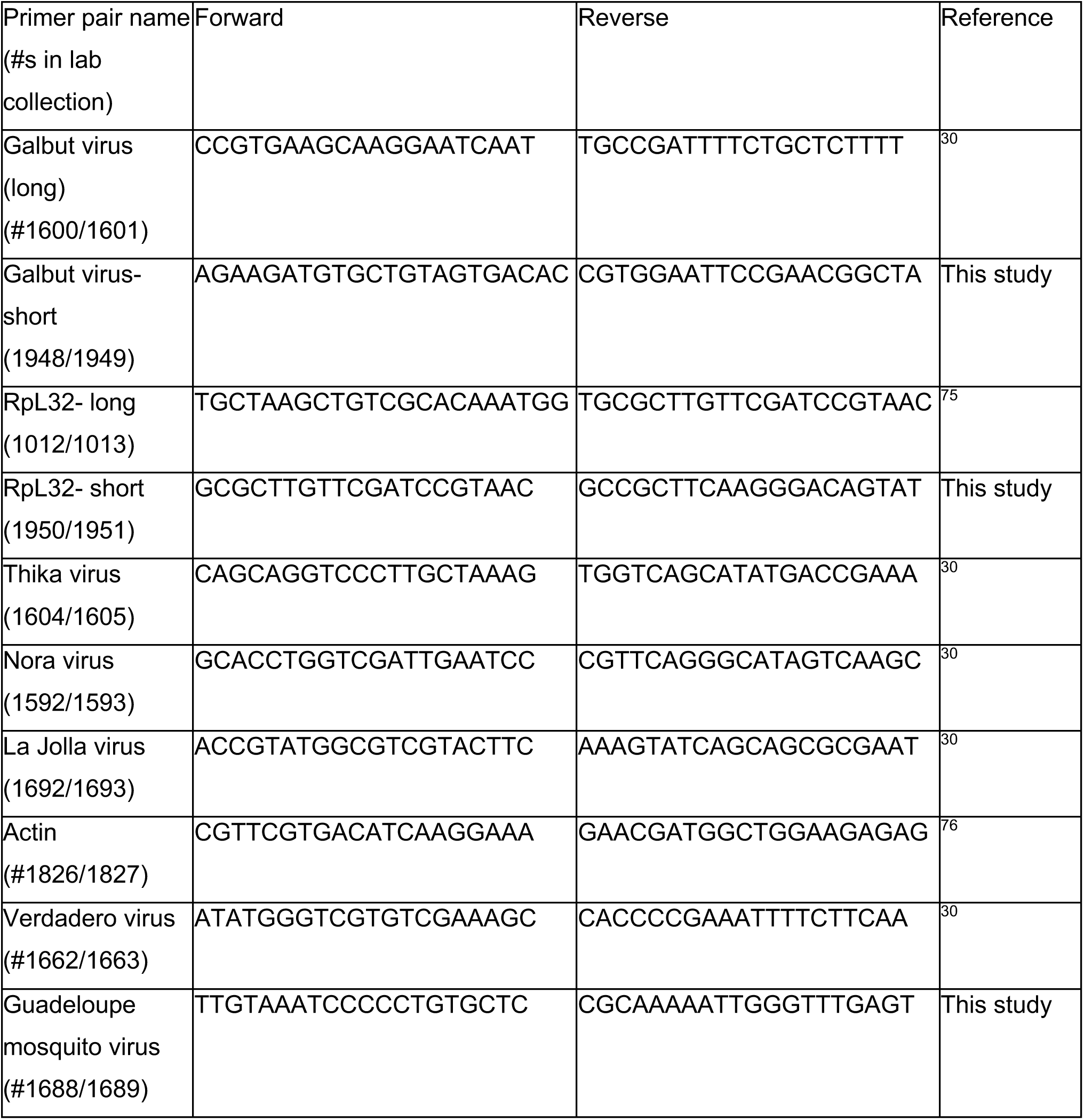
Primer sequences used in this study.

### NGS Library Preparation

Museum collection samples were prepared for sequencing using the Kapa Biosystems Kapa RNA Hyper Prep kit following manufacturer recommendations with modification of the fragmentation/priming step. Since the RNA was already highly fragmented, samples were minimally incubated at 65°C for 1 minute during the fragmentation/priming step (Roche, 08098093702). Due to low starting concentrations, 12 cycles were used for library amplification. FoCo-17 fly RNA was used as a positive control and water was used as a negative control for library preparation and sequencing. Controls were sequenced on a separate run to avoid index hopping from the positive control sample^77^. Sample pooling was determined using High Sensitivity DNA Qubit reagents and final library quality control was done using an Agilent D1000 HS Tapestation and KapaQuant reagents following manufacturers recommendations (Roche, 07960140001). Museum samples were sequenced on three NextSeq 500 runs; one high output 75 cycle run and two mid output 150 cycle runs. All museum specimens were run on the first NextSeq run, runs two and three consisted of samples that we wanted to increase coverage of near complete virus sequences. Experimentally dried samples were sequenced on a high output 75 cycle run. The two positive control samples were sequenced on a 300 cycle MiSeq run. Additional fresh frozen samples were sequenced on a NextSeq mid output 150 cycle kit.

## Data Analysis

*Experimental Collections:* All statistical analysis and data visualization, unless otherwise noted, was done using R/RStudio. Data cleaning and data visualization were done using the tidyverse package^78^. All scripts are available in the GitHub repository linked below. A p-value of α = 0.05 or below was used as our significance threshold. RNA Concentration and Length: Four individual multiple linear regression models (MLRs) were constructed to test for statistical significance in RNA concentration and length between dried and frozen flies and mosquitoes over 72- and 52-weeks. The predictor time was treated as a continuous variable and the predictor storage type was converted to factor. A balanced one-way analysis of variance (ANOVA) was used to test for significance using the car package^79^. The performance package was used to test model assumptions^80^. RT-qPCR: Raw qPCR data files were cleaned and compiled using the SangerTools R package and a custom R script^81^. Two-tailed Students T-tests from the rstatix package were used to determine statistical significance of dried and frozen Ct values^82^. P-values were adjusted for multiple hypothesis testing.

*Identification of virus sequences in old samples:* We used our lab’s previously described metagenomic classification pipeline (https://github.com/stenglein-lab/taxonomy_pipeline) to identify and validate virus sequences in NGS datasets^30^. In brief, adapters and low quality reads were trimmed using cutadapt 3.5^83^. Fastqc was used to assess post-collapsed read quality 0.11.9^84^. Host reads were removed using bowtie2 2.4.5 and the remaining reads were assembled using spades 3.15.4^85,86^. Virus sequences were identified using BLASTn 2.12.0 and a BLASTx search using diamond 2.0.14 was used to identify novel sequences^87,88^. Draft sequences were validated by remapping of trimmed reads using bwa mem aligner version 0.7.17^89^. Final sequences were submitted to the NCBI nucleotide database and raw NGS data to the NCBI sequence read archive repository.

This metagenomic classification pipeline was also used to taxonomically assign non-host contigs and unassembled reads. Counts assigned to individual taxa were normalized to the number of quality- and adapter-trimmed reads.

*Molecular species identification using cytochrome oxidase subunit 1 competitive mapping.* The sequence of this mitochondrial-encoded gene is commonly used to distinguish related species^90^. We used the *D. melanogaster* CO1 coding sequence from NC_024511 as a BLASTN query to the NCBI nucleotide database to identify other drosophilid CO1 sequences^91^. We restricted results to sequences from Drosophilidae and to alignments with >90% query coverage and E-value <1x10^-50^. We used cd-hit-est to collapse sequences that shared >99% pairwise nucleotide identity to create a set of 545 representative CO1 sequences^92^. We mapped adapter and quality trimmed reads to these sequences using bowtie2 v.2.4.4 in end-to-end mode and retained alignments with a mapping quality ≥ 20.

*Maximum Likelihood Trees:* All available galbut virus and sigma virus sequences from individual flies (and not pools) with collection date metadata were downloaded from GenBank. Sequences in the Puslinch virus phylogeny include all available L protein sequences in the NCBI RefSeq protein database from the *Peribunyaviridae* family. Sequences in the sobemo-like virus tree include RdRp protein sequences from viral genomes identified by a BLASTN search of the NCBI nucleotide database. Alignments were generated using MAFTT v7.490 with default parameters. IqTree v1.0 (http://iqtree.cibiv.univie.ac.at) was used to generate maximum likelihood trees^93^. The following parameters were used on the web interface version of IqTree: substitution model-auto, Free-rate Heterogeneity-yes, bootstrap analysis-ultrafast with all other options set to default. FigTree was used for tree visualization (http://tree.bio.ed.ac.uk/software/figtree/). Trees were midpoint rooted.

*Analysis of strandedness of virus-mapping reads.* To tabulate the numbers of positive and negative strand reads mapping to virus sequences, we mapped quality- and adapter-trimmed reads to assembled virus sequences using the bwa mem aligner version 0.7.17^89^. The orientation of each mapped read was determined from the 0x10 bit flag of the resulting sam format output.

For all analyses of virus- and host -mapping reads, values were summarized and visualized using the tidyverse R packages^78^. Wilcoxon tests were used to statistically evaluate differences between groups using the rstatix R package. In all cases values were determined to be non-normally distributed by Shapiro-Wilk test^82^. *P*-values were adjusted for multiple testing using the Holm–Bonferroni method and were indicated on plots using the ggpubr R package using the following significance levels: ns: p > 0.05; *: p <= 0.05; **: p <= 0.01; ***: p <= 0.001; ****: p <= 0.0001^94^. Analyes were implemented as nextflow workflows available at: https://github.com/LKeene/Old_Collections_Figures/^95^.

*Analysis of ribosomal rRNA-mapping reads.* Adapter- and quality-trimmed reads from *D. melanogaster* datasets were mapped to the 18S and 28S rRNA sequences from Anger et al (chains B5 and A2 from^54^) using bwa mem aligner version 0.7.17^89^. Per-base, per-strand coverage was calculated using the BamToCov tool v2.7.0^96^. The *D. melanogaster* 80S ribosome structure was visualized using ChimeraX v.1.7.1^97^. Bases were binned into one of four categories (paired, non-paired, higher-order structure, absent from structure) by feeding this structure (protein data bank accession 4V6W) into RNApdbee 2.0^55^.

*Analysis of non-ribosomal RNA types. A*dapter- and quality-trimmed reads from *D. melanogaster* datasets were mapped to the *D. melanogaster* genome version BDGP6.32^98^ using bwa mem aligner version 0.7.17^89^. Mapping output and genome annotation were input into ALFA v1.1.1 to tabulate the fractions of reads mapping to each of the annotated genomic biotypes^57^.

*Analysis of mismatch signatures of chemical damage.* Trimmed reads from *D. melanogaster* datasets were mapped to rRNA as above. Mismatches of bases with a basecall quality score >30 were tabulated.

*Competitive mapping to ribosomal internal transcribed spacer sequences.* We collected sequences between the 18S and 5.8S rRNA genes for the four museum species. Sequences correspond to Genbank accessions: *D. melanogaster*: NR_133558.1:2861-722; *D. simulans*: NGVV02000020.1:5755-5071; *D. putrida*: AF184042.1:153-611; *S. pallida*: JAECXP010000218.1:7393-7904. We mapped adapter and quality trimmed reads to these sequences using bowtie2 v.2.4.4 in end-to-end mode and retained alignments with a mapping quality ≥ 30.

## ACKNOWLEDGEMENTS

The authors would like to acknowledge the museum institutions and staff for their willingness to share specimens for this work: Robin Thomson, University of Minnesota Insect Collection; Timothy McCabe, New York State Museum; Chris Paradise, The Davidson College Entomology Collection; Matt Gimmel, Santa Barbara Museum of Natural History; Camiel Doorenweerd, University of Hawaii Insect Museum; Jayme Sones, The Centre for Biodiversity and Genomics at the University of Guelph; Laura Porturas, The Frost Entomology Center at Pennsylvania State University; Tommy McElrath, Illinois Natural History Survey. The authors would also like to thank Dr. Brian Foy for providing *Ae. aegypti* mosquitoes, and Dr. Rebekah Kading for the supplies to pin and store dried insects. We thank Darren Obbard, Dan Sloan, and Jeff Wilusz for helpful discussions.

## DATA AVAILABILITY

All metadata, RT-qPCR raw data, plots, and analysis and visualization code can be found at: https://github.com/LKeene/Old_Collections_Figures. Raw sequence data has been uploaded to the NCBI SRA database; assembled viral sequences have been deposited in GenBank; both can be found under NCBI BioProject accession PRJNA1034757.

## REFERENCES

1. Leonardi, M. et al. Evolutionary Patterns and Processes: Lessons from Ancient DNA. Syst Biol 66, syw059 (2016).

2. Orlando, L. & Cooper, A. Using ancient DNA to understand evolutionary and ecological processes. Annu Rev Ecol Evol Syst 45, 573–598 (2014).

3. Green, R. E. et al. A draft sequence of the neandertal genome. Science (1979) 328, 710– 722 (2010).

4. Duchêne, S., Ho, S. Y. W., Carmichael, A. G., Holmes, E. C. & Poinar, H. The Recovery, Interpretation and Use of Ancient Pathogen Genomes. Current Biology 30, R1215–R1231 (2020).

5. Ng, T. F. F. et al. Preservation of viral genomes in 700-y-old caribou feces from a subarctic ice patch. Proc Natl Acad Sci U S A 111, 16842–7 (2014).

6. Fromm, B. et al. Ancient microRNA profiles of 14,300-yr-old canid samples confirm taxonomic origin and provide glimpses into tissue-specific gene regulation from the Pleistocene. RNA 27, 324–334 (2021).

7. Faria, N. R. et al. The early spread and epidemic ignition of HIV-1 in human populations. Science (1979) 346, 56–61 (2014).

8. Taubenberger, J. K., Reid, A. H., Krafft, A. E., Bijwaard, K. E. & Fanning, T. G. Initial Genetic Characterization of the 1918 “Spanish” Influenza Virus. Science (1979) 275, 1793–1796 (1997).

9. Belyi, V. A., Levine, A. J. & Skalka, A. M. Sequences from Ancestral Single-Stranded DNA Viruses in Vertebrate Genomes: the Parvoviridae and Circoviridae Are More than 40 to 50 Million Years Old. J Virol 84, 12458–12462 (2010).

10. Ávila-Arcos, M. C. et al. One hundred twenty years of koala retrovirus evolution determined from museum Skins. Mol Biol Evol 30, 299–304 (2013).

11. Enard, D. & Petrov, D. A. Ancient RNA virus epidemics through the lens of recent adaptation in human genomes. Philosophical Transactions of the Royal Society B: Biological Sciences 375, 20190575 (2020).

12. Yang, E. et al. Decay rates of human mRNAs: correlation with functional characteristics and sequence attributes. Genome Res 13, 1863–72 (2003).

13. Fordyce, S. L., Kampmann, M.-L., van Doorn, N. L. & Gilbert, M. T. P. Long-term RNA persistence in postmortem contexts. Investig Genet 4, 7 (2013).

14. Smith, O. et al. A complete ancient RNA genome: Identification, reconstruction and evolutionary history of archaeological Barley Stripe Mosaic Virus. Sci Rep 4, 4003 (2014).

15. Peyambari, M., Warner, S., Stoler, N., Rainer, D. & Roossinck, M. J. A 1,000-Year-Old RNA Virus. J Virol 93, e01188–18 (2019).

16. Mármol-Sánchez, E. et al. Historical RNA expression profiles from the extinct Tasmanian tiger. Genome Res 33, 1299–1316 (2023).

17. Cobb, N. S. et al. Assessment of North American arthropod collections: prospects and challenges for addressing biodiversity research. PeerJ 7, e8086 (2019).

18. Watts, P. C., Thompson, D. J., Allen, K. A. & Kemp, S. J. How useful is DNA extracted from the legs of archived insects for microsatellite-based population genetic analyses? J Insect Conserv 11, 195–198 (2007).

19. Heintzman, P. D., Elias, S. A., Moore, K., Paszkiewicz, K. & Barnes, I. Characterizing DNA preservation in degraded specimens of Amara alpina (Carabidae: Coleoptera). Mol Ecol Resour 14, 606–615 (2014).

20. Goldstein, P. Z. & Desalle, R. Calibrating phylogenetic species formation in a threatened insect using DNA from historical specimens. Mol Ecol 12, 1993–1998 (2003).

21. Tin, M. M.-Y., Economo, E. P. & Mikheyev, A. S. Sequencing Degraded DNA from Non-Destructively Sampled Museum Specimens for RAD-Tagging and Low-Coverage Shotgun Phylogenetics. PLoS One 9, e96793 (2014).

22. Lalonde, M. M. L. & Marcus, J. M. How old can we go? Evaluating the age limit for effective DNA recovery from historical insect specimens. Syst Entomol 45, 505–515 (2020).

23. Gilbert, M. T. P., Moore, W., Melchior, L. & Worobey, M. DNA Extraction from Dry Museum Beetles without Conferring External Morphological Damage. PLoS One 2, e272 (2007).

24. Shpak, M., Ghanavi, H. R., Lange, J. D., Pool, J. E. & Stensmyr, M. C. Genomes from historical Drosophila melanogaster specimens illuminate adaptive and demographic changes across more than 200 years of evolution. PLoS Biol 21, e3002333 (2023).

25. Webster, C. L., Longdon, B., Lewis, S. H. & Obbard, D. J. Twenty-Five New Viruses Associated with the Drosophilidae (Diptera). Evolutionary Bioinformatics 12s2, EBO.S39454 (2016).

26. Webster, C. L. et al. The Discovery, Distribution, and Evolution of Viruses Associated with Drosophila melanogaster. PLoS Biol 13, e1002210 (2015).

27. Yamaguchi, M. & Yoshida, H. Drosophila as a Model Organism. in Advances in Experimental Medicine and Biology vol. 1076 1–10 (Springer New York LLC, 2018).

28. Beckingham, K. M., Armstrong, J. D., Texada, M. J., Munjaal, R. & Baker, D. A. Drosophila melanogaster--the model organism of choice for the complex biology of multi-cellular organisms. Gravit Space Biol Bull 18, 17–29 (2005).

29. Longdon, B., Wilfert, L. & Jiggins, F. M. The Sigma Viruses of Drosophila from: Rhabdoviruses: Molecular Taxonomy, Evolution, Genomics, Ecology, Host-Vector Interactions, Cytopathology and Control. (Caister Academic Press, U.K., 2012).

30. Cross, S. T. et al. Partitiviruses Infecting Drosophila melanogaster and Aedes aegypti Exhibit Efficient Biparental Vertical Transmission. J Virol 94, (2020).

31. Vera-Maloof, F. Z., Saavedra-Rodriguez, K., Elizondo-Quiroga, A. E., Lozano-Fuentes, S. & Black Iv, W. C. Coevolution of the Ile1,016 and Cys1,534 Mutations in the Voltage Gated Sodium Channel Gene of Aedes aegypti in Mexico. PLoS Negl Trop Dis 9, e0004263 (2015).

32. Ortiz-Baez, A. S., Shi, M., Hoffmann, A. A. & Holmes, E. C. RNA virome diversity and Wolbachia infection in individual Drosophila simulans flies. J Gen Virol 102, 001639 (2021).

33. Habayeb, M. S., Ekengren, S. K. & Hultmark, D. Nora virus, a persistent vitus in Drosophila, defines a new picorna-like virus family. Journal of General Virology 87, 3045– 3051 (2006).

34. Shi, C. et al. Stable distinct core eukaryotic viromes in different mosquito species from Guadeloupe, using single mosquito viral metagenomics. Microbiome 7, 121 (2019).

35. Santos, D., Ribeiro, G. C., Cabral, A. D. & Sperança, M. A. A non-destructive enzymatic method to extract DNA from arthropod specimens: Implications for morphological and molecular studies. PLoS One 13, e0192200 (2018).

36. Sturtevant, A. H. A New Species Closely Resembling *Drosophila Melanogaster*. Psyche (Camb Mass) 26, 153–155 (1919).

37. Sturtevant, A. H. Notes on North American Drosophilidae with Descriptions of Twenty-Three New Species. Ann Entomol Soc Am 9, 323–343 (1916).

38. Cross, S. T. et al. Co-Infection Patterns in Individual Ixodes scapularis Ticks Reveal Associations between Viral, Eukaryotic and Bacterial Microorganisms. Viruses 10, 388 (2018).

39. Hoskins, R. A. et al. The Release 6 reference sequence of the Drosophila melanogaster genome. Genome Res 25, 445–58 (2015).

40. Kim, B. Y. et al. Single-fly genome assemblies fill major phylogenomic gaps across the Drosophilidae Tree of Life. PLoS Biol 22, e3002697 (2024).

41. Balique, F., Colson, P. & Raoult, D. Tobacco mosaic virus in cigarettes and saliva of smokers. Journal of Clinical Virology 55, 374–376 (2012).

42. Kircher, M., Sawyer, S. & Meyer, M. Double indexing overcomes inaccuracies in multiplex sequencing on the Illumina platform. Nucleic Acids Res 40, e3 (2012).

43. Rambaut, A., Lam, T. T., Max Carvalho, L. & Pybus, O. G. Exploring the temporal structure of heterochronous sequences using TempEst (formerly Path-O-Gen). Virus Evol 2, vew007 (2016).

44. Aiewsakun, P. & Katzourakis, A. Time-Dependent Rate Phenomenon in Viruses. J Virol 90, 7184–7195 (2016).

45. Ghafari, M., Simmonds, P., Pybus, O. G. & Katzourakis, A. A mechanistic evolutionary model explains the time-dependent pattern of substitution rates in viruses. Current Biology 31, 4689–4696.e5 (2021).

46. Drummond, A., Pybus, O. G. & Rambaut, A. Inference of viral evolutionary rates from molecular sequences. Adv Parasitol 54, 331–58 (2003).

47. Duchêne, S., Holmes, E. C. & Ho, S. Y. W. Analyses of evolutionary dynamics in viruses are hindered by a time-dependent bias in rate estimates. Proceedings of the Royal Society B: Biological Sciences 281, 20140732 (2014).

48. Cross, S. T. et al. Galbut Virus Infection Minimally Influences Drosophila melanogaster Fitness Traits in a Strain and Sex-Dependent Manner. Viruses 15, 539 (2023).

49. Wallace, M. A. et al. The discovery, distribution, and diversity of DNA viruses associated with *Drosophila melanogaster* in Europe. Virus Evol 7, veab031 (2021).

50. Lowen, A. C. It’s in the mix: Reassortment of segmented viral genomes. PLoS Pathog 14, e1007200 (2018).

51. Parkhomchuk, D. et al. Transcriptome analysis by strand-specific sequencing of complementary DNA. Nucleic Acids Res 37, e123 (2009).

52. Nibert, M. L. et al. Taxonomic reorganization of family Partitiviridae and other recent progress in partitivirus research. Virus Res 188, 128–41 (2014).

53. Robicheau, B. M., Susko, E., Harrigan, A. M. & Snyder, M. Ribosomal RNA genes contribute to the formation of pseudogenes and junk DNA in the human genome. Genome Biol Evol 9, 380–397 (2017).

54. Anger, A. M. et al. Structures of the human and Drosophila 80S ribosome. Nature 497, 80–5 (2013).

55. Zok, T. et al. RNApdbee 2.0: multifunctional tool for RNA structure annotation. Nucleic Acids Res 46, W30–W35 (2018).

56. Staple, D. W. & Butcher, S. E. Pseudoknots: RNA structures with diverse functions. PLoS Biol 3, e213 (2005).

57. Bahin, M. et al. ALFA: annotation landscape for aligned reads. BMC Genomics 20, 250 (2019).

58. Sharp, S. J., Schaack, J., Cooley, L., Burke, D. J. & Söll, D. Structure and transcription of eukaryotic tRNA genes. CRC Crit Rev Biochem 19, 107–44 (1985).

59. Zhang, J. Recognition of the tRNA structure: Everything everywhere but not all at once. Cell Chem Biol 31, 36–52 (2024).

60. Dabney, J., Meyer, M. & Pääbo, S. Ancient DNA damage. Cold Spring Harb Perspect Biol 5, a012567 (2013).

61. Shibutani, S., Takeshita, M. & Grollman, A. P. Insertion of specific bases during DNA synthesis past the oxidation-damaged base 8-oxodG. Nature 349, 431–4 (1991).

62. Sloan, D. B., Broz, A. K., Sharbrough, J. & Wu, Z. Detecting Rare Mutations and DNA Damage with Sequencing-Based Methods. Trends Biotechnol 36, 729–740 (2018).

63. Rhee, Y., Valentine, M. R. & Termini, J. Oxidative base damage in RNA detected by reverse transcriptase. Nucleic Acids Res 23, 3275–82 (1995).

64. Sinha, R. et al. Index switching causes ‘spreading-of-signal’ among multiplexed samples in Illumina HiSeq 4000 DNA sequencing. doi:10.1101/125724.

65. Valentine, M. R. & Termini, J. Kinetics of formation of hypoxanthine containing base pairs by HIV-RT: RNA template effects on the base substitution frequencies. Nucleic Acids Res 29, 1191–9 (2001).

66. Zhang, K., Hodge, J., Chatterjee, A., Moon, T. S. & Parker, K. M. Duplex structure of double-stranded RNA provides stability against hydrolysis relative to single-stranded RNA. Environ Sci Technol 55, 8045–8053 (2021).

67. Sandoval-Velasco, M. et al. Three-dimensional genome architecture persists in a 52,000-year-old woolly mammoth skin sample. Cell 187, 3541–3562.e51 (2024).

68. Petrzik, K. Evolutionary forces at work in partitiviruses. Virus Genes 55, 563–573 (2019).

69. Skoglund, P. et al. Separating endogenous ancient DNA from modern day contamination in a Siberian Neandertal. Proc Natl Acad Sci U S A 111, 2229–2234 (2014).

70. Laurence, M., Hatzis, C. & Brash, D. E. Common contaminants in next-generation sequencing that hinder discovery of low-abundance microbes. PLoS One 9, e97876 (2014).

71. Naccache, S. N. et al. The perils of pathogen discovery: origin of a novel parvovirus-like hybrid genome traced to nucleic acid extraction spin columns. J Virol 87, 11966–77 (2013).

72. Kjær, K. H. et al. A 2-million-year-old ecosystem in Greenland uncovered by environmental DNA. Nature 612, 283–291 (2022).

73. Krogmann, L. & Holstein, J. Chapter 18 Preserving and Specimen Handling: Insects and Other Invertebrates. vol. 8 (ABC TAXA, 2010).

74. Foster, W. A. Colonization and Maintenance of Mosquitoes in the Laboratory. in *Pathology*, Vector Studies, and Culture 103–151 (Elsevier, 1980). doi:10.1016/B978-0-12-426102-0.50009-9.

75. Cao, C., Magwire, M. M., Bayer, F. & Jiggins, F. M. A Polymorphism in the Processing Body Component Ge-1 Controls Resistance to a Naturally Occurring Rhabdovirus in Drosophila. PLoS Pathog 12, e1005387 (2016).

76. Dzaki, N., Ramli, K. N., Azlan, A., Ishak, I. H. & Azzam, G. Evaluation of reference genes at different developmental stages for quantitative real-time PCR in Aedes aegypti. Sci Rep 7, 43618 (2017).

77. van der Valk, T., Vezzi, F., Ormestad, M., Dalén, L. & Guschanski, K. Index hopping on the Illumina HiseqX platform and its consequences for ancient DNA studies. Mol Ecol Resour 20, 1171–1181 (2020).

78. Wickham, H. et al. Welcome to the Tidyverse. J Open Source Softw 4, 1686 (2019).

79. Fox, J. & Weisberg, S. An R Companion to Applied Regression. (Sage, 2019).

80. Lüdecke, D., Ben-Shachar, M., Patil, I., Waggoner, P. & Makowski, D. performance: An R Package for Assessment, Comparison and Testing of Statistical Models. J Open Source Softw 6, 3139 (2021).

81. Laldin, A. & Hutson, G. SangerTools: Tools for Population Health Management Analytics. (2022).

82. Kassambara, A. rstatix: Pipe-Friendly Framework for Basic Statistical Tests. https://rpkgs.datanovia.com/rstatix/ (2023).

83. Martin, M. Cutadapt Removes Adapter Sequences From High-Throughput Sequencing Reads. EMBnet J 17, 10–12 (2011).

84. Andrews, S., et al. FastQC: a quality control tool for high throughput sequence data. https://qubeshub.org/resources/fastqc (2012).

85. Langmead, B. & Salzberg, S. L. Fast gapped-read alignment with Bowtie 2. Nat Methods 9, 357–359 (2012).

86. Prjibelski, A., Antipov, D., Meleshko, D., Lapidus, A. & Korobeynikov, A. Using SPAdes De Novo Assembler. Curr Protoc Bioinformatics 70, e102 (2020).

87. Buchfink, B., Reuter, K. & Drost, H. G. Sensitive protein alignments at tree-of-life scale using DIAMOND. Nat Methods 18, 366–368 (2021).

88. Camacho, C. et al. BLAST+: architecture and applications. BMC Bioinformatics 10, 421 (2009).

89. Li, H. & Durbin, R. Fast and accurate long-read alignment with Burrows-Wheeler transform. Bioinformatics 26, 589–595 (2010).

90. Hebert, P. D. N., Ratnasingham, S. & deWaard, J. R. Barcoding animal life: cytochrome c oxidase subunit 1 divergences among closely related species. Proc Biol Sci 270 Suppl 1, S96–9 (2003).

91. Morgulis, A. et al. Database indexing for production MegaBLAST searches. Bioinformatics 24, 1757–64 (2008).

92. Fu, L., Niu, B., Zhu, Z., Wu, S. & Li, W. CD-HIT: accelerated for clustering the next-generation sequencing data. Bioinformatics 28, 3150–2 (2012).

93. Nguyen, L. T., Schmidt, H. A., Von Haeseler, A. & Minh, B. Q. IQ-TREE: A fast and effective stochastic algorithm for estimating maximum-likelihood phylogenies. Mol Biol Evol 32, 268–274 (2015).

94. Kassambara, A. ggpubr: ‘ggplot2’ Based Publication Ready Plots. R package version 0.6.0. https://rpkgs.datanovia.com/ggpubr/ (2023).

95. Zhang, J. et al. International Cancer Genome Consortium Data Portal--a one-stop shop for cancer genomics data. Database (Oxford) 2011, bar026 (2011).

96. Birolo, G. & Telatin, A. BamToCov: An efficient toolkit for sequence coverage calculations. Bioinformatics 38, 2617–2618 (2022).

97. Meng, E. C. et al. UCSF ChimeraX: Tools for structure building and analysis. Protein Sci 32, e4792 (2023).

98. Hoskins, R. A. et al. The Release 6 reference sequence of the Drosophila melanogaster genome. Genome Res 25, 445–458 (2015).

