## Supplemental Figures for "Sequencing RNA from old, dried specimens reveals past viromes and properties of long-surviving RNA"

***Viral metagenomics of 100-year-old museum specimens highlights the long-term stability of RNA***

**Supplemental Figures**

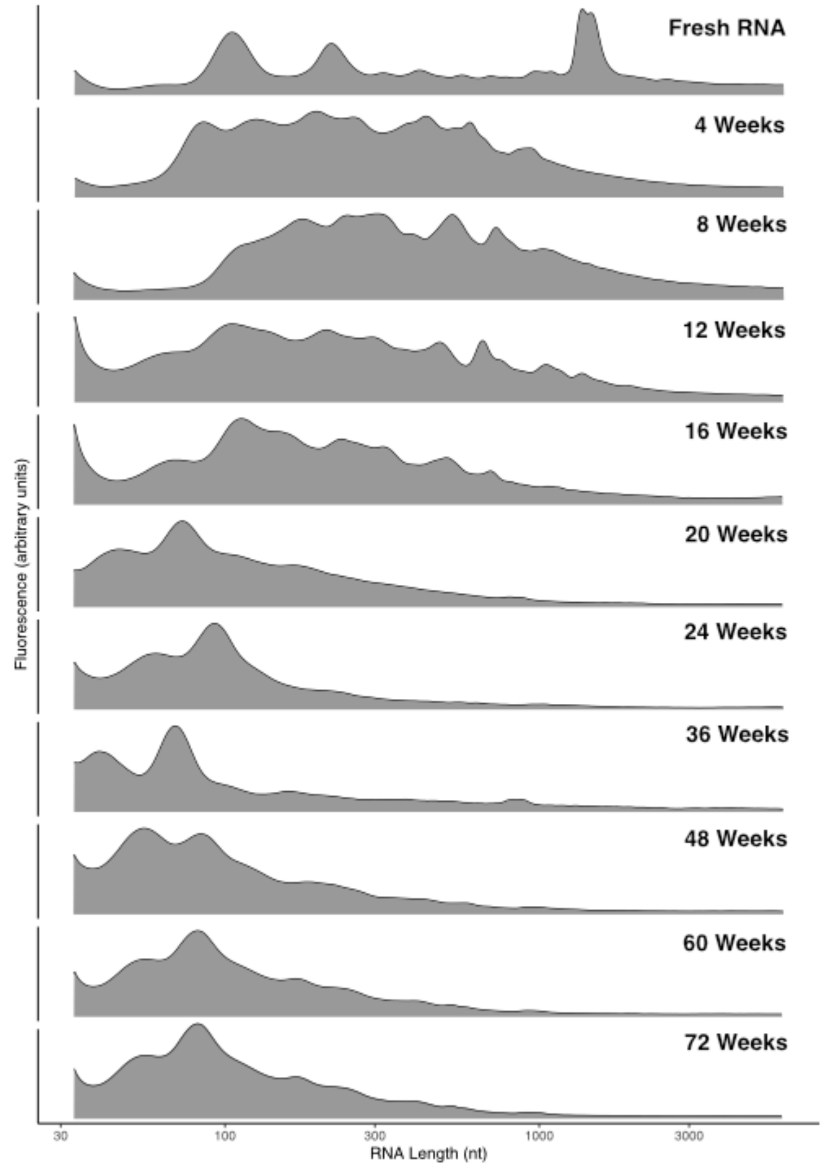


**Supplemental Figure 1: RNA grew increasingly fragmented in experimentally dried flies.** The RNA length of representative sample from each time point of dried *D. melanogaster* are shown. An Agilent Tapestation was used to measure RNA length distributions. Lengths <33 nt, corresponding to the Tapestation lower marker internal control, are not plotted.

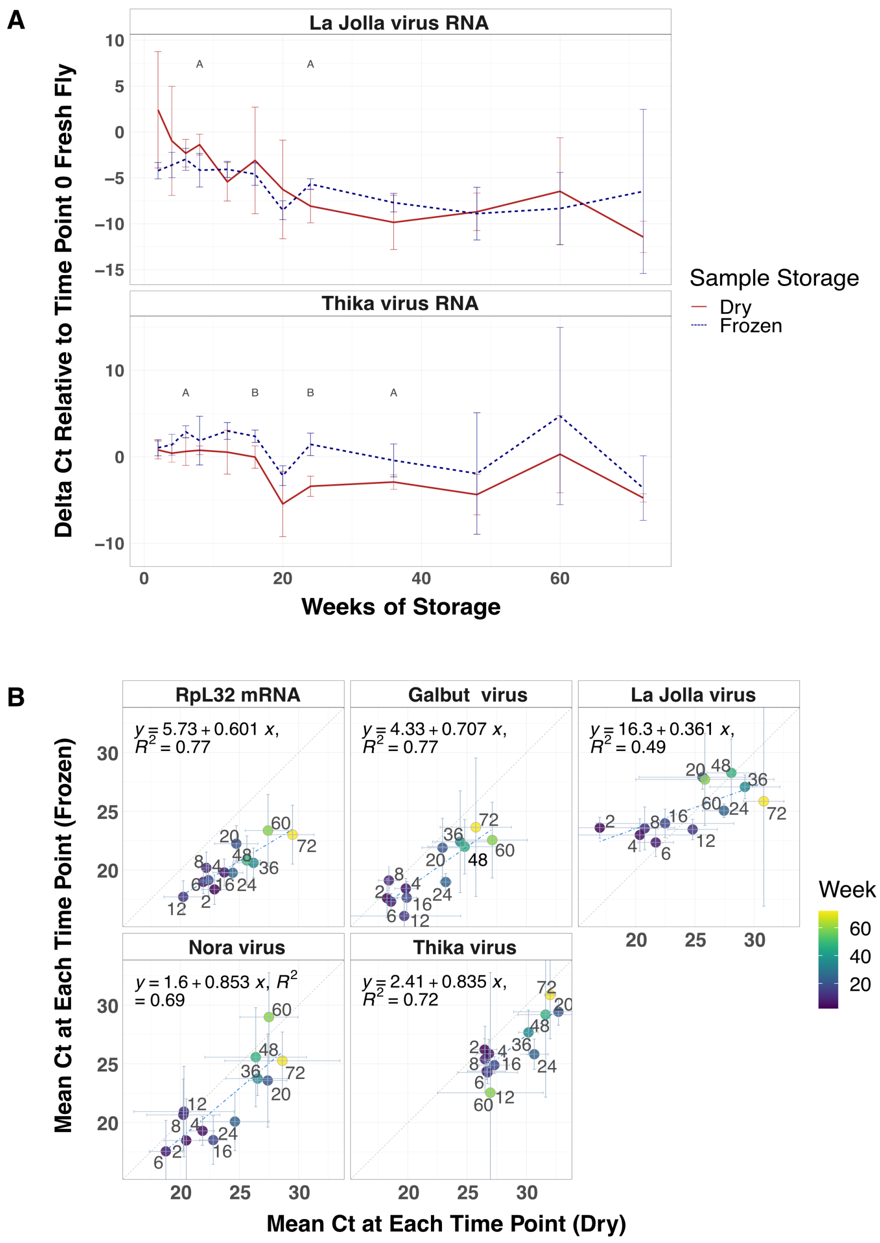


**Supplementary Figure 2: Viral RNA and host mRNA persisted in fly specimens over 72 weeks.** (A) Average difference between La Jolla virus and Thika virus RNA levels relative to levels in fresh *D. melanogaster*. A two-tailed t-test was used to determine the statistical significance between dried and frozen means at every time point (A < 0.05, B < 0.001). (B) Comparison of mean Ct values at each time point for each RNA target. Linear regression lines are shown as is the diagonal. Error bars represent mean plus and minus the standard deviation. N = 3 male and 3 female at each time point.

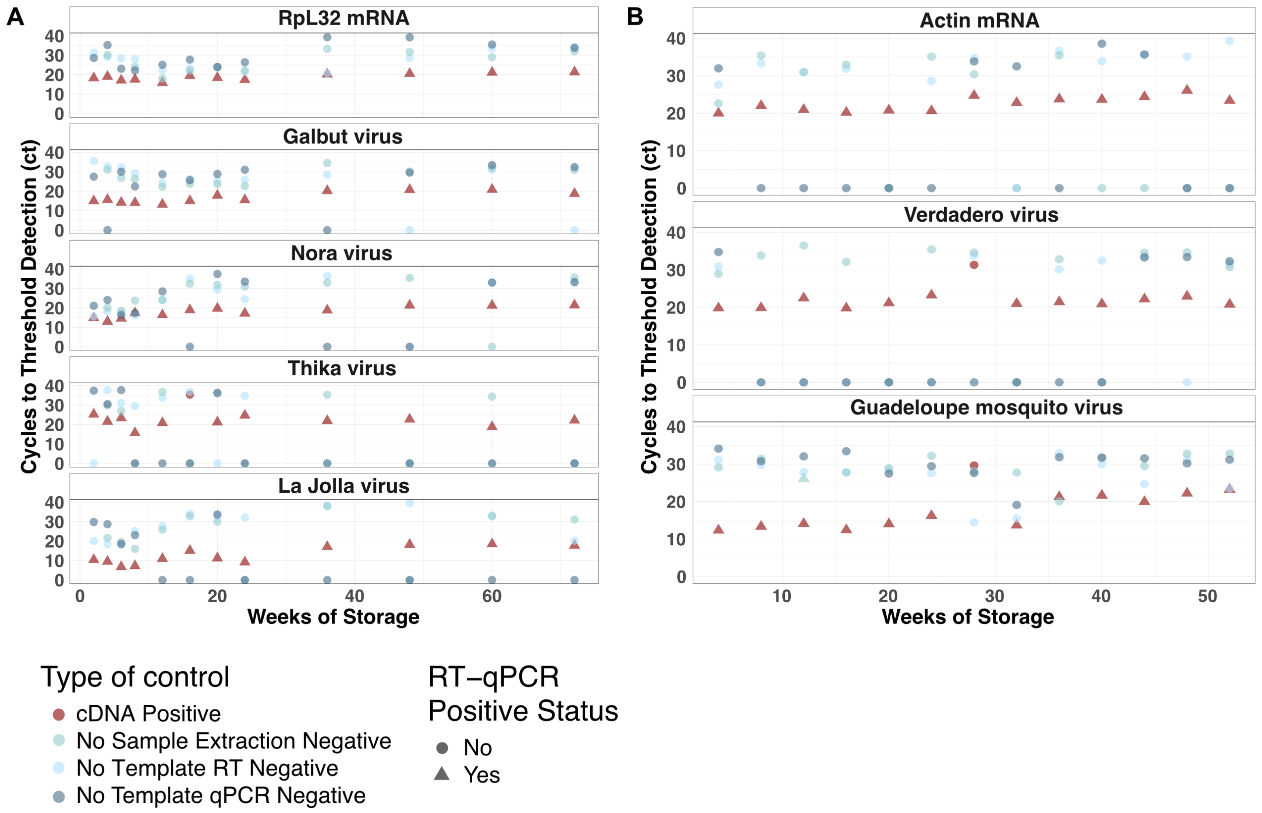


**Supplemental Figure 3: Positive and negative controls in experimentally dried flies and mosquitoes showed no evidence of cross-contamination at extraction or RT-qPCR steps.** Detection and threshold cycle for the indicated RNA targets are shown for experimentally dried and frozen flies (A) and mosquitoes (B). One positive and 3 negative controls were used at each time point: a cDNA positive control, a no-sample extraction blank control, a no-template RT control, and a no-sample qPCR control. Positive calls required the sample's melting temperature to be within one degree of the cDNA positive control's. Undetermined Cts (no amplification) were set to zero. Samples from negative control samples with called Cts (likely from primer dimer products) were confirmed negative by analysis of product melting temperatures and agarose gel electrophoresis. Color indicates control type and symbols indicate whether samples were called positive or negative.

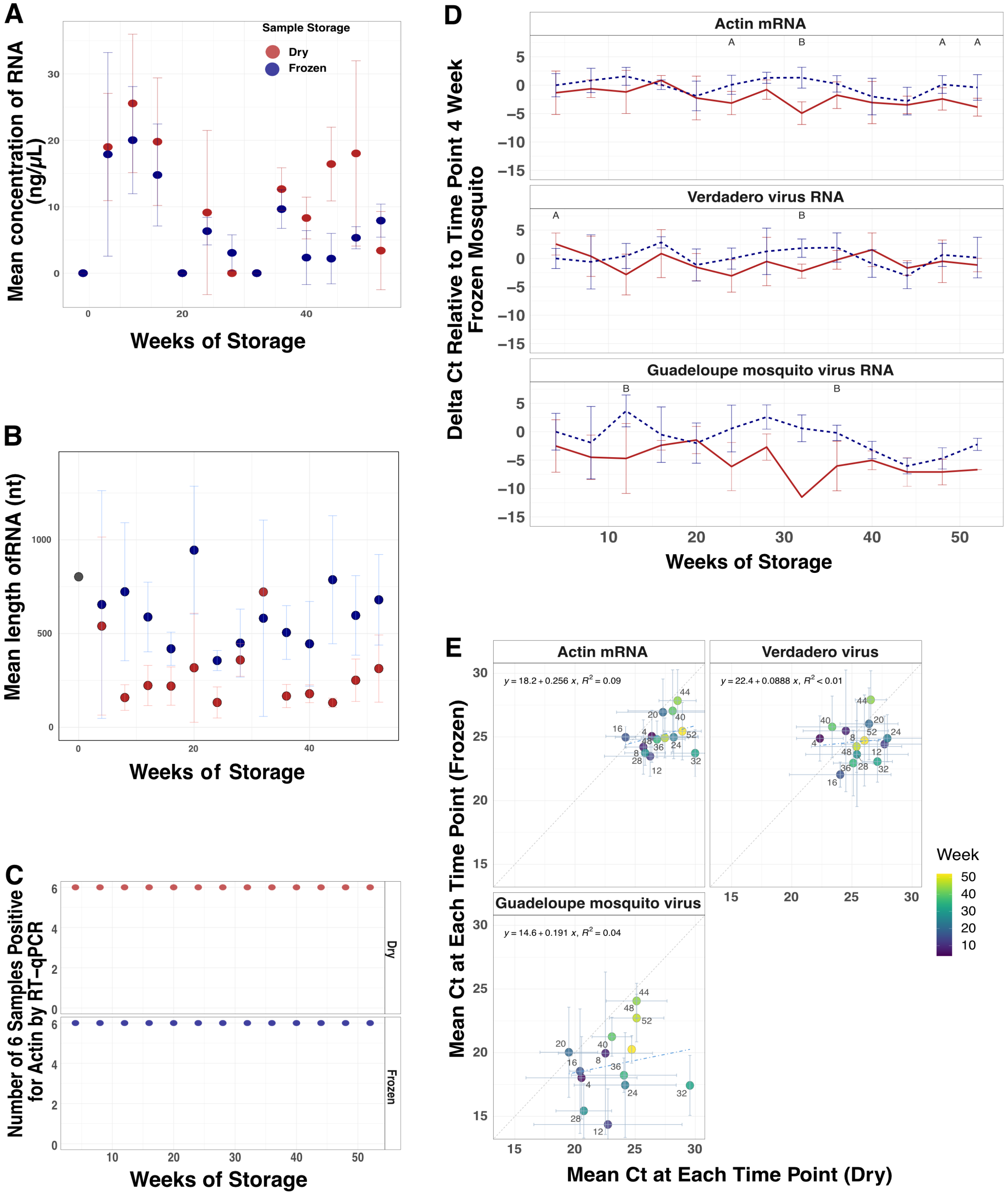
A
Sample Storage
Dry
Frozen
Mean concentration of RNA (ng/μL)
Weeks of Storage
B
Mean length ofRNA (nt)
Weeks of Storage
C
Number of 6 Samples Positive for Actin by RT-qPCR
Dry
Frozen
Weeks of Storage
D
Delta Ct Relative to Time Point 4 Week Frozen Mosquito
Actin mRNA
Verdadero virus RNA
Guadeloupe mosquito virus RNA
Weeks of Storage
E
Actin mRNA
y = 18.2 + 0.256 x, R² = 0.09
Verdadero virus
y = 22.4 + 0.0888 x, R² < 0.01
Guadeloupe mosquito virus
y = 14.6 + 0.191 x, R² = 0.04
Mean Ct at Each Time Point (Frozen)
Mean Ct at Each Time Point (Dry)
Week

**Supplemental Figure 4: Viral RNA and host mRNA persisted in dried mosquito specimens over 52 weeks.** (A) Average RNA concentration and (B) RNA length of RNA from dried and frozen mosquitoes over 52 weeks. N = 3 female dried or frozen *Ae. aegypti* at each time. (C) Number of samples positive of 3 males and 3 females for actin mRNA via RT-qPCR in dried and frozen specimens over 52 weeks. (D) Average difference of actin mRNA, verdadero virus and Guadeloupe mosquito virus RNA levels relative to time point 4 weeks frozen *Ae. aegypti*. A two-tailed t-test was used to determine the statistical significance between dried and frozen means at every time point (A < 0.05, B < 0.001). (E) Comparison of mean Ct values at each time point for each RNA target. Linear regression lines are shown as is the diagonal. Error bars represent mean plus and minus the standard deviation. N = 3 male and 3 female.

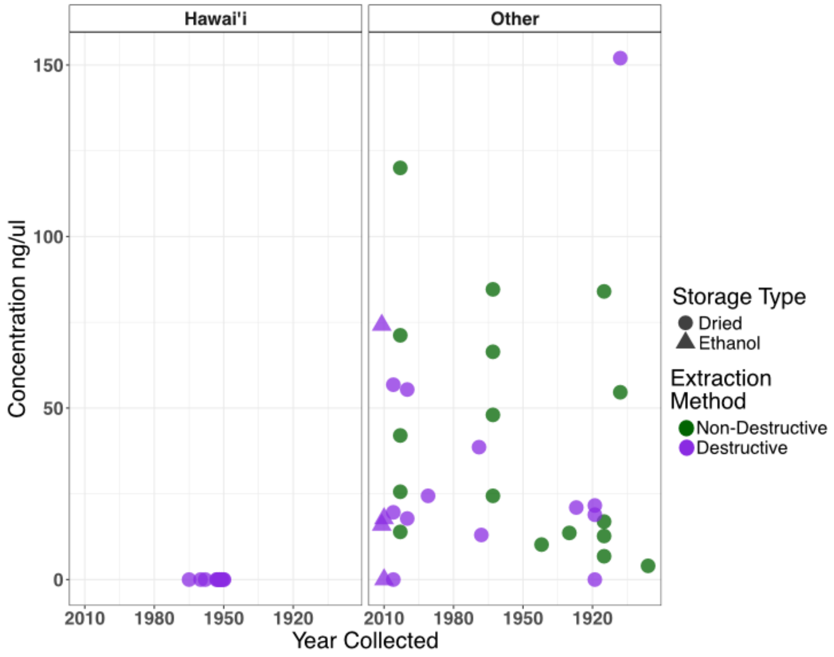


**Supplemental Figure 5: RNA yield from Hawai'ian specimens was low.** RNA concentrations of samples from Hawai'i or any other location. Purple = destructive RNA extraction and green = non-destructive RNA extraction. Shape indicates whether the sample was dried or stored in ethanol.

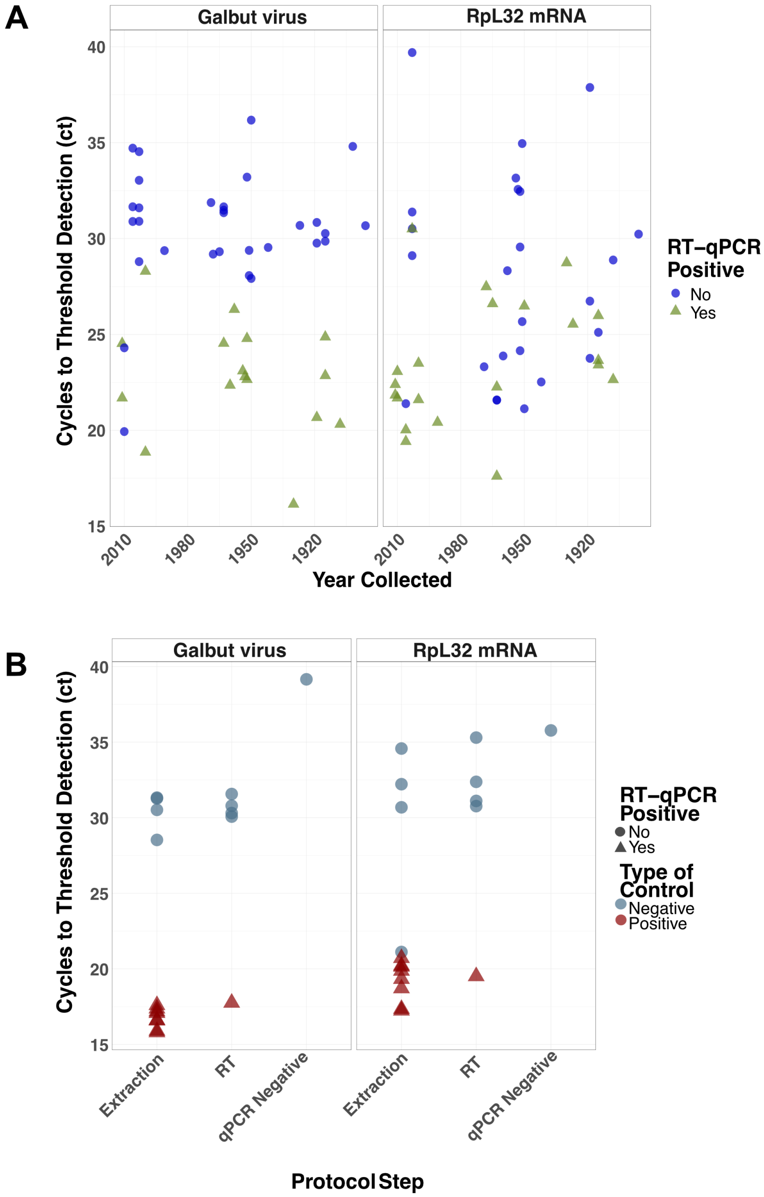


**Supplemental Figure 6: Screening of Museum specimens and controls for galbut virus and RpL32 mRNA.** (A) RT-qPCR results for museum specimens and (B) controls screened for galbut virus and RpL32 mRNA using short-range primers. Positive calls required that the melting temperature be within one degree of the cDNA positive control and the presence of an appropriately sized band on an agarose gel to distinguish legitimate positives from primer dimers. Undetermined Cts were set to zero. Two extraction positive (fresh FoCo-17 fly) and one negative (extraction blank) control was used for each set of extractions. Four RT no template negative controls were also used to increase the chance of detecting contamination.

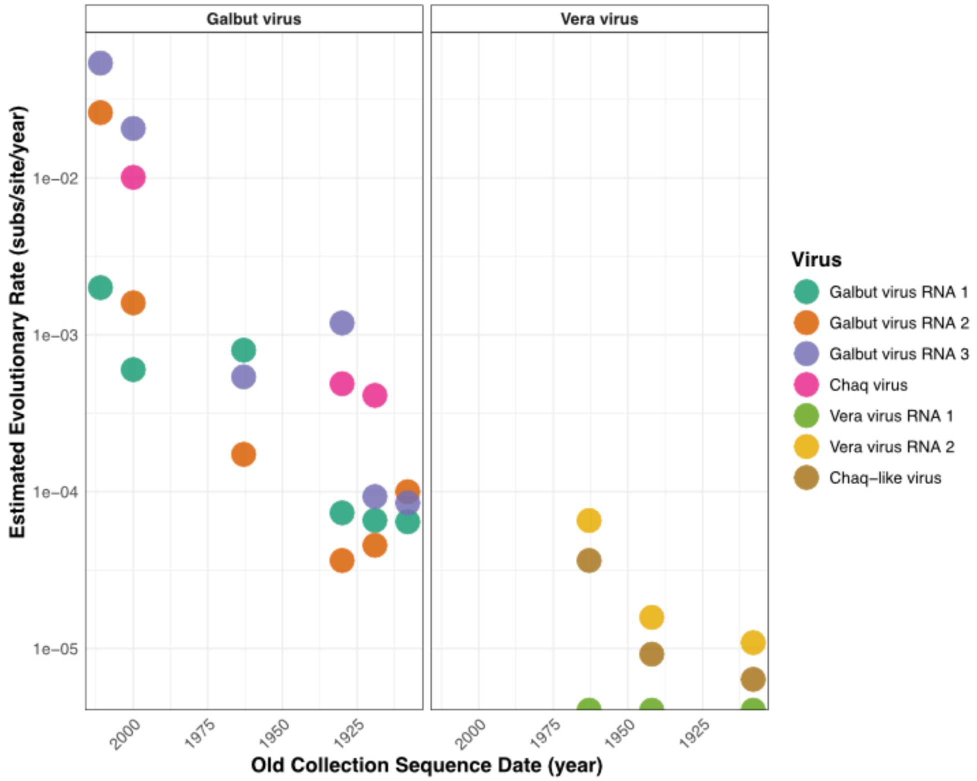


**Supplemental Figure 7: The estimated evolutionary rate of galbut virus and vera virus from sequences recovered from museum specimens.** Rates were calculated as number of substitutions per nucleotide per year relative to the most closely related sequence available in GenBank with a known collection year. Samples without a suitably close contemporary sequence were removed from this analysis. Color indicates virus. Vera virus RNA 1 sequences are 100% identical to their closest available sequence.

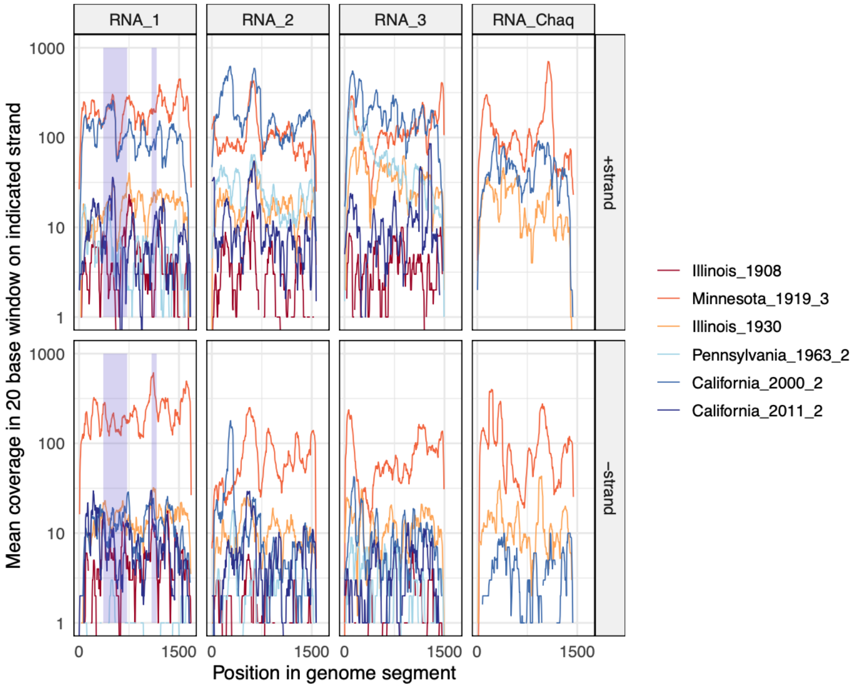


**Supplemental Figure 8: Galbut virus coverage derived from both strands of all segments**. Coverage levels of +strand and –strand mapping reads across galbut virus segments are plotted. The position of the long and short amplicons targeting RNA 1 are shaded in purple (**Supplemental Table 2**).

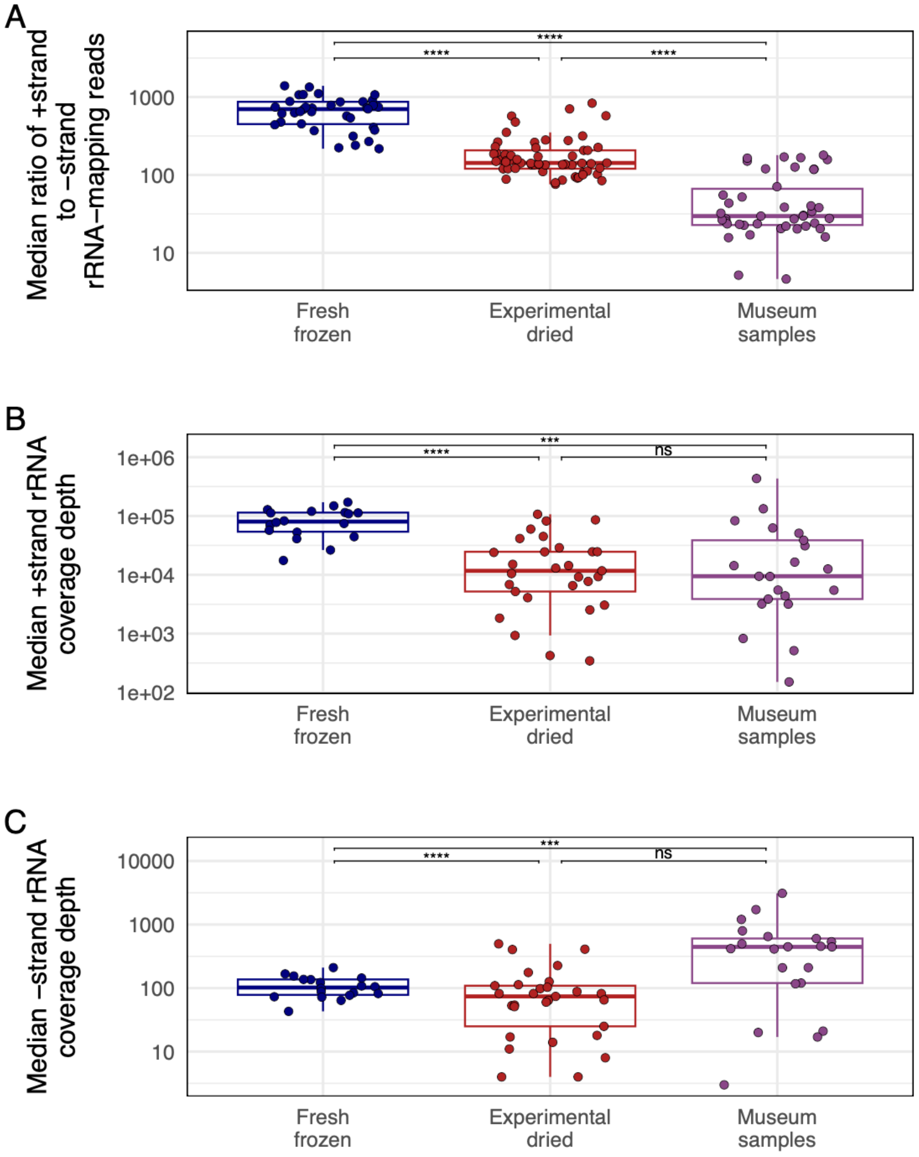


**Supplemental Figure 9: Strand ratios of surviving ribosomal RNA were consistent with preferential survival of dsRNA in old samples**. (A) The ratio of +strand to –strand coverage of rRNA-mapping reads in *D. melanogaster* datasets. Each point represents the median ratio from individual-fly datasets. Adjusted *p*-values significance levels from Wilcoxon test are indicated. (B) The depth of coverage of +strand rRNA-mapping reads. (C) The depth of coverage of –strand rRNA mapping reads.

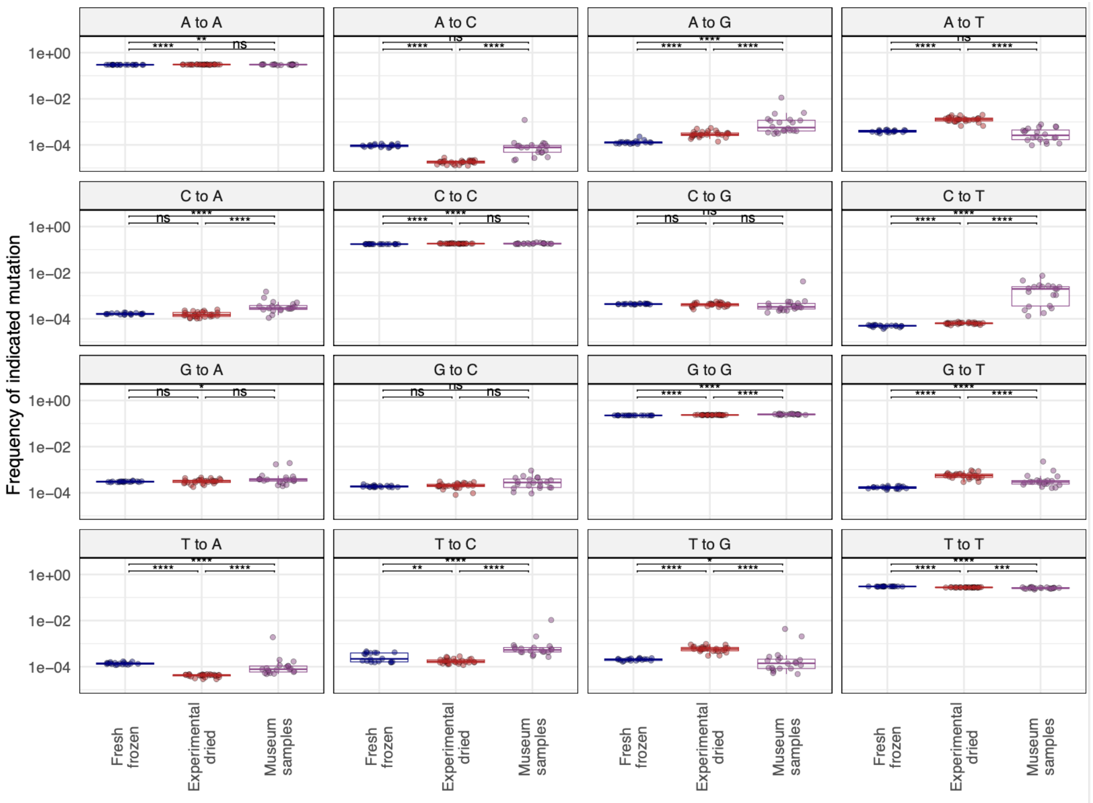


**Supplemental Figure 10: Old RNA was chemically damaged and exhibits mismatch patterns consistent with cytosine and adenine deamination.** Mismatches in rRNA-mapping reads from *D. melanogaster* datasets were quantified and the frequency of each mismatch type is plotted. Each point represents a dataset from an individual fly. Adjusted *p*-values significance levels from Wilcoxon test are indicated.

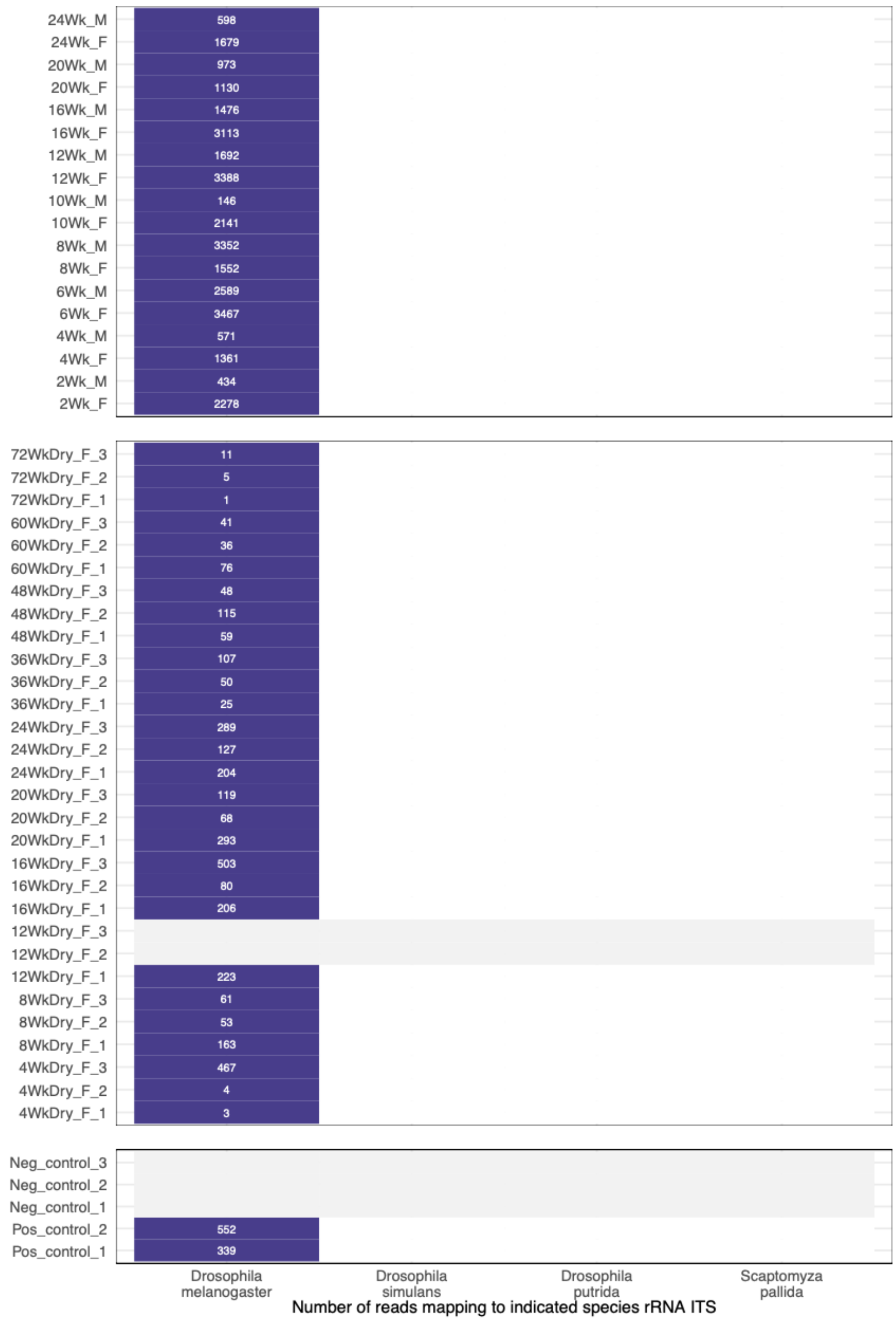


**Supplemental Figure 11: Host-mapping reads exhibited species specificity.** The number of reads mapping to the indicated species rRNA internal transcribed spacer 1 (ITS) sequence for each fresh-frozen, experimentally dried, or control dataset is shown. Datasets with no ITS-mapping reads, including negative control water libraries, are represented with grey.
